# Competition between hematopoietic stem and progenitor cells controls hematopoietic stem cell compartment size

**DOI:** 10.1101/2021.12.12.472293

**Authors:** Runfeng Miao, Harim Chun, Ana Cordeiro Gomes, Jungmin Choi, João P. Pereira

## Abstract

Cellular competition for limiting hematopoietic factors is a physiologically regulated but poorly understood process. Here, we studied this phenomenon by hampering hematopoietic progenitor access to Leptin receptor^+^ mesenchymal stem/progenitor cells (MSPCs) and endothelial cells (ECs). We show that HSC numbers increased by 2-fold when multipotent and lineage-restricted progenitors failed to respond to CXCL12 produced by MSPCs and ECs. HSCs were qualitatively normal, and HSC expansion only occurred when early hematopoietic progenitors but not differentiated hematopoietic cells lacked CXCR4. Furthermore, the MSPC and EC transcriptomic heterogeneity was remarkably stable, suggesting that it is impervious to dramatic changes in hematopoietic progenitor interactions. Instead, HSC expansion was caused by increased availability of membrane-bound stem cell factor (mSCF) on MSPCs and ECs due to reduced consumption by cKit-expressing hematopoietic progenitors. These studies revealed an intricate homeostatic balance between HSCs and proximal hematopoietic progenitors regulated by cell competition for limited amounts of mSCF.

## Introduction

In adult mammals, hematopoietic stem cells (HSCs) rely on a combination of key paracrine signals provided by specialized microenvironments in the bone marrow and by the liver. While HSCs can access long-range acting signals, such as hepatocyte-produced Thrombopoietin (Decker et al., 2018), presumably anywhere, other signals such as Stem Cell Factor (SCF, or Kit ligand encoded by *Kitl*) and CXCL12 are preferentially accessed in specialized niches formed predominantly by perisinusoidal mesenchymal stem/progenitor cells (MSPCs) and endothelial cells (ECs) in the bone marrow (Asada et al., 2017; Ding and Morrison, 2013; Ding et al., 2012; Greenbaum et al., 2013; Mendez-Ferrer et al., 2010; Morrison and Scadden, 2014; Omatsu et al., 2010; Xu et al., 2018). Although historically defined as hematopoietic stem cell niches, more recent studies demonstrated that these niches are also critical for the development of B-lymphoid lineage cells due to the fact that MSPCs and ECs are also exclusive cellular sources of IL7, a key lymphopoietic cytokine (Cordeiro Gomes et al., 2016; Fistonich et al., 2018; Lim et al., 2017; Zehentmeier and Pereira, 2019). Besides IL7, CXCL12, and SCF, MSPCs and ECs also express several key hematopoietic cytokines with well-defined roles in myeloid and lymphoid-lineage cell differentiation, such as FLT3L, MCSF, IL34, IL15, GCSF among others, as revealed by single-cell RNA sequencing of non-hematopoietic bone marrow cell populations (Baccin et al., 2020; Baryawno et al., 2019; Tikhonova et al., 2019). Importantly, common myeloid and lymphoid progenitors, and megakaryocyte and erythroid progenitors depend on SCF produced by Lepr+ MSPCs, whereas macrophage and dendritic cell precursors (MDPs) and monocytes depend on MCSF produced by ECs in bone marrow (Comazzetto et al., 2019; Shen et al., 2021; Zhang et al., 2021). Combined, these studies led us to propose that MSPCs and at least some ECs are not only required for the long-term maintenance of HSCs but also form appropriate environments for the development of most, if not all, lymphoid and myeloid cell lineages (Miao et al., 2020). Terminally differentiated hematopoietic cell subsets, namely macrophages, megakaryocytes, and regulatory T cells (Tregs), can in turn relay signals such as adenosine, TGFβ, and PF4 back to the HSC, or alter HSC niche activity through changes in CXCL12 production, and control the size of the HSC compartment under homeostatic conditions (Bruns et al., 2014; Chow et al., 2011; Hirata et al., 2018; Zhao et al., 2014).

While some hematopoietic cytokines produced by MSPCs and ECs act in cell lineage-restricted manners (e.g., MCSF in monocyte/macrophage development; IL7 in lymphoid lineage development, etc.), other signals such as SCF are shared by multiple hematopoietic progenitor subsets. This type of cellular organization in which MSPCs and ECs harbor a constellation of hematopoietic cells and nurture distinct cell lineages raises the possibility that competition between HSCs and downstream progenitors for common and limiting resources could control HSCs and hematopoietic progenitors under homeostasis and during perturbations. However, arguments have been made against this possibility. Specifically, MSPCs and ECs outnumber HSCs by more than 10 fold, indicating that many putative HSC niches may remain “vacant” (Shimoto et al., 2017; Wei and Frenette, 2018). But, when taking into account not only the number of HSCs but also of downstream progenitors (e.g., CMPs, CLPs, MEPs, GMPs, etc.), the stoichiometry between the number of niche cells and of hematopoietic stem and progenitor cells is lower than one.

In this study, we tested whether cellular competition between HSCs and hematopoietic progenitors for HSC niche factors could control the HSC compartment size. HSCs and hematopoietic progenitor and differentiated cells utilize the CXCR4/CXCL12 pathway for bone marrow homing, and for access to the marrow parenchyma where they are retained via adhesive interactions with CXCL12-producing MSPCs and ECs. By examining the impact of CXCR4 conditional deletion at multiple stages of hematopoietic cell development, we found that when MPPs are deficient in CXCR4, the HSC compartment size increased by ~ 2-fold without any measurable loss of HSC fitness. This increase occurred without any major changes in the MSPC and EC transcriptome nor in transcriptional heterogeneity of the non-hematopoietic compartment. Surprisingly, HSC expansion was entirely controlled by excess membrane-bound SCF (mSCF) on MSPCs and ECs due to its reduced consumption by cKit-expressing hematopoietic progenitor cells. These studies provide new insights into the fine-balance between HSCs and downstream progenitors regulated by a poorly understood phenomenon of cellular competition in the hematopoietic organ.

## Results

### Control of HSC compartment size requires hematopoietic progenitor interactions with niche cells

In prior studies, we noted that HSCs and MPPs could be found in close proximity to each other and to the same bone marrow niche cell (Cordeiro Gomes et al., 2016), suggesting that individual niche cells support a variety of hematopoietic stem and progenitor cells. Consistent with this possibility, recent studies demonstrated that besides HSCs, lymphoid, myeloid, and erythroid precursors require SCF produced by Lepr+ MSPCs (Comazzetto et al., 2019; Shen et al., 2021; Zhang et al., 2021). These studies led us to ask if competition between HSCs and downstream hematopoietic progenitors for factors locally produced by niche cells could control the HSC compartment size. Most signals produced by niche cells act in a short-range manner, and access to such signals is dependent on localization cues of which CXCL12 is the most abundantly produced. Thus, we analyzed mice in which HSCs express CXCR4 and respond to its ligand CXCL12 while MPPs and downstream hematopoietic progenitor and differentiated cells lack CXCR4 via conditional deletion using *Flk2*-driven Cre recombinase (Fig. 1A). The *Flk2*-cre genetic approach has been described to efficiently target MPPs and downstream hematopoietic cells leaving HSCs unchanged (Boyer et al., 2011). We found that phenotypic long-term HSC numbers were increased by ~ 2-fold in the bone marrow, while short-term HSCs and MPP2 and MPP3 cell populations (Pietras et al., 2015) remained unchanged (Fig. 1B and C, and Fig. S1A and B). The number of lymphoid-primed MPP4 cells was dramatically reduced in bone marrow (Fig. 1C and S1A and B) and increased in the spleen (Fig. 1D), consistent with a critical role for CXCR4 in HSC and hematopoietic progenitor cell retention in bone marrow (Beck et al., 2014; Cordeiro Gomes et al., 2016). We also observed increased numbers of phenotypic HSCs in the spleen (Fig. 1D), suggesting an overall increase in medullary and extra-medullary HSCs. The number of LSKs in bone marrow was unchanged (Fig. 1E). Other myeloid and lymphoid progenitors and differentiated immune cells were also significantly reduced in the bone marrow (Fig. S1A and B), as expected (Cordeiro Gomes et al., 2016). To assess if the number of functional HSCs was increased, we analyzed their ability to re-populate the HSC compartment of lethally irradiated recipient mice. In a setting of mixed bone marrow transplantation with 50% bone marrow cells from CXCR4 conditionally deficient mice (*Flk2*-cre^+^ *Cxcr4^fl/fl^*, from here on referred to as cKO) and 50% bone marrow cells from wild-type (WT) C57BL6/NCI mice distinguished by CD45 isoforms (CD45.1 and CD45.2), the mixed chimerism in the HSC compartment of cKO:WT chimeric mice was significantly higher than that in control WT:WT mixed chimeras 16 weeks after hematopoietic reconstitution (Fig. 1F). In sharp contrast, the mixed chimerism in MPP4s and in differentiated hematopoietic cells was dramatically reduced in cKO:WT chimeric mice (Fig. S1C), as expected (Cordeiro Gomes et al., 2016). Collectively, these data revealed that phenotypic HSCs become numerically increased when MPP4 and downstream hematopoietic progenitors lack CXCR4.

**Figure 1.**
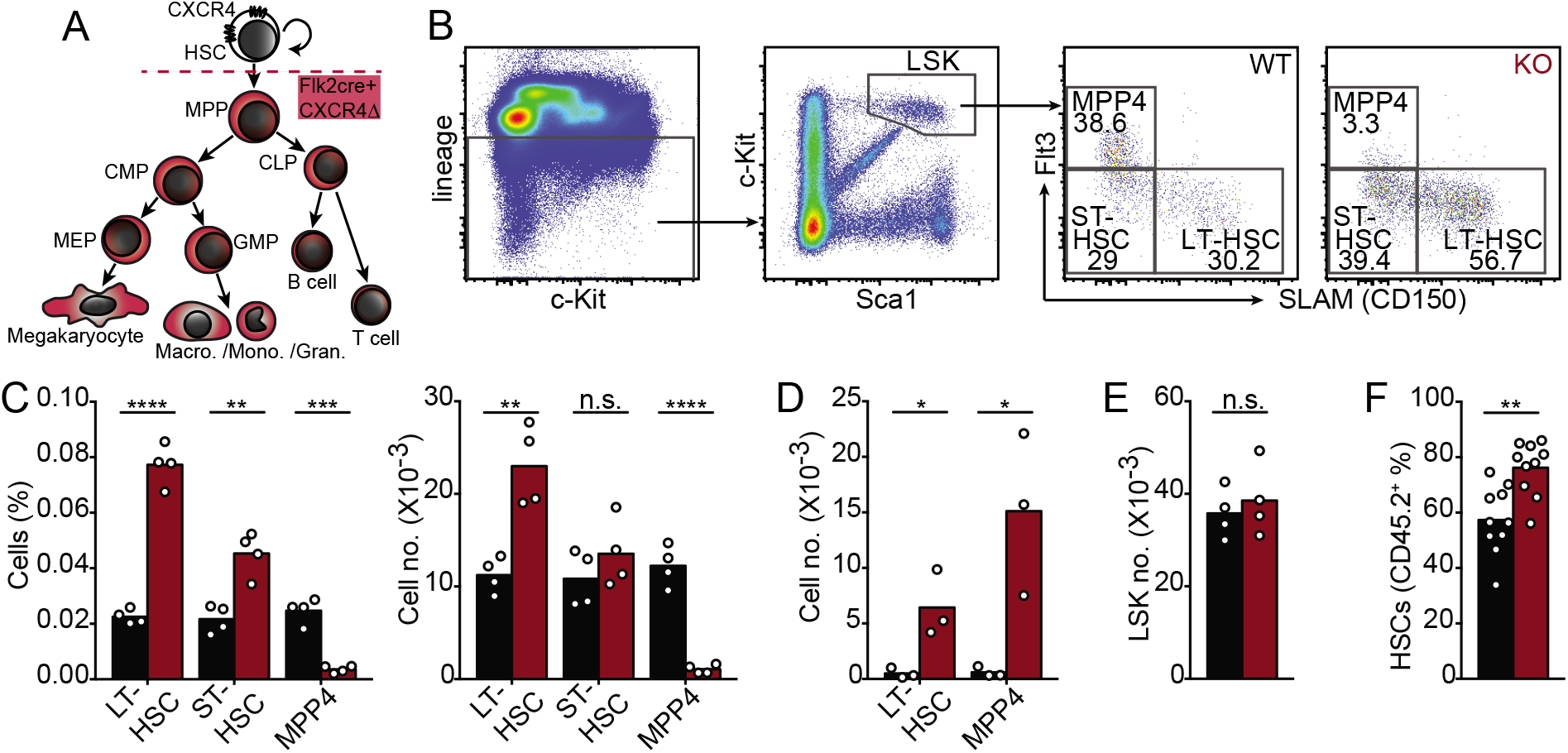
Increased HSC numbers in mice with CXCR4-deficient MPPs. (A) Schematic representation of CXCR4 deletion in hematopoietic multipotent progenitor cells with a *Flk2*-cre transgene. (B) Gating strategy for LSKs, LT-HSC, ST-HSC, and MPP4 cells. (C and D) LT-HSC, ST-HSC, and MPP4 cell numbers in bone marrow (C) and in spleen (D). (E) LSK cell number in bone marrow. In panels C-E, cells were collected from one femur and tibia, or from spleen, of *Flk2-cre.Cxcr4^fl/+^* (CTR, black) and *Flk2-cre.Cxcr4^fl/fl^* mice (cKO, red). (F) HSC chimerism in bone marrow of mice reconstituted with 50% CD45.2^+^ CTR or cKO bone marrow cells mixed with 50% CD45.1^+^ wild-type bone marrow cells. Data in all panels are representative of two or more experiments. Bars indicate average, circles depict individual mice. *, P < 0.05; **, P < 0.01; *** P < 0.001; and ****, P < 0.0001 by unpaired Student’s *t* test.

### HSC quiescence and self-renewal are independent of CXCR4-mediated hematopoietic progenitor localization

Increased numbers of phenotypic HSCs in cKO mice suggested increased entry into cell cycle, and possibly reduced quiescence. To test this, we analyzed intracellular levels of the nuclear protein associated with cell proliferation Ki67 in phenotypic HSCs of both mice by flow cytometry. Although we found an increased frequency of Ki67^+^ HSCs in cKO mice (Fig. 2A), this did not result in reduced numbers of Ki67^-^ quiescent HSCs (Fig. 2B). Furthermore, when challenging mice with the myelosuppressive agent 5-Fluorouracil (5-FU, weekly treatment i.p. at 150 mg/Kg), cKO mice were equally resistant to 5-FU treatment as WT littermate controls (Fig. 2C), which contrasted sharply with increased susceptibility to 5-FU when HSCs and downstream hematopoietic cells lack CXCR4 (Sugiyama et al., 2006). To determine if phenotypic HSCs were functionally normal we examined their capacity for long-term self-renewal and multilineage differentiation in vivo. In transplantation experiments of 50% mixed bone marrow chimeras (50% cKO or littermate and 50% CD45.1+ C57BL6), the mixed chimerism of HSCs, B lymphocytes and granulocytes from cKO bone marrow remained stable after 16 weeks of primary bone marrow transplantation followed by another 16 weeks of secondary transplantation (Fig. 2D). These results showed that HSCs in cKO mice were functionally equivalent to HSCs from control littermate mice. To specifically determine if phenotypic HSCs were functionally normal, we sorted HSCs from cKO mice or control littermate (30 phenotypic HSCs) and transplanted into lethally irradiated C57BL6 (CD45.1+) recipient mice, followed by secondary transplantation into new cohorts of C57BL6 (CD45.1+) recipient mice. The mixed chimerism in the phenotypic HSC, CD19+ B cell, and Gr1+ granulocyte compartments was determined 16 weeks after the primary and secondary transplant. HSCs from WT and cKO mice were equivalent in their ability to self-renew and to differentiate into lymphoid and myeloid-lineage cells upon primary and secondary transplantation (Fig. 2E-G). Consistent with these observations, HSCs isolated from cKO and WT mice displayed similar expression of CD150 or CD41 on the cell surface (data not shown), markers whose increased expression is associated with myeloid differentiation bias (Challen et al., 2010; Gekas and Graf, 2013). Combined, these studies show a numerical increase in phenotypic and functional HSCs in cKO mice without loss of HSC quiescence, self-renewal, or differentiation bias.

**Figure 2.**
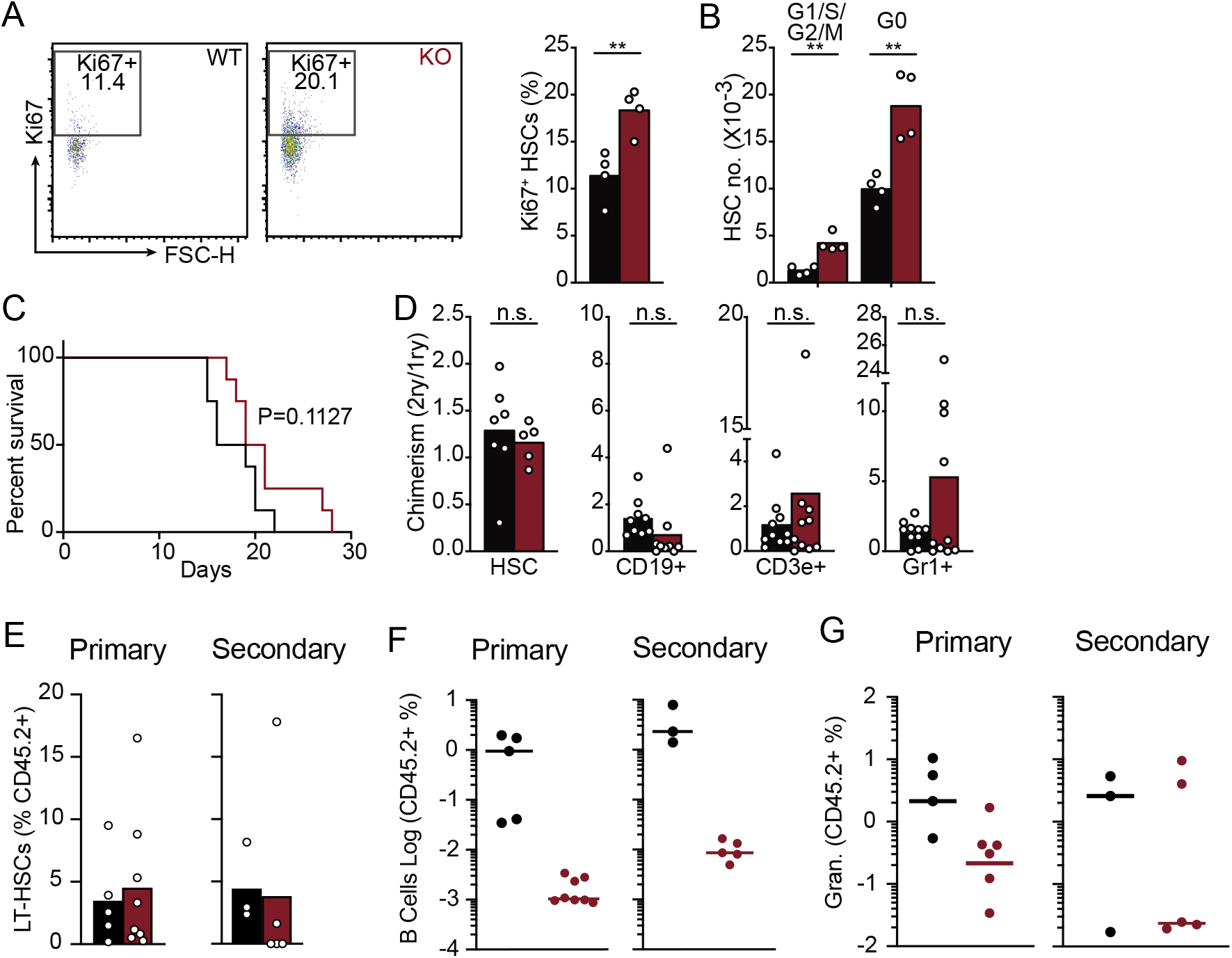
Measurement of HSC quiescence and self-renewal in mice with CXCR4-deficient MPPs. (A and B) HSC cell cycle status. (A) Gating strategy and cell frequency; (B) Enumeration of HSCs in G0 (Ki67^-^) and in G1/S/G2M (Ki67^+^) in bone marrow. (C) Kaplan-Meyer survival plot of mice following weekly 5-FU treatment (i.p., 150mg/kg, n=8 per group). (A-C) data obtained from *Flk2-cre.Cxcr4^fl/+^* (black) and *Flk2-cre.Cxcr4^fl/fl^* (red) mice (D) Serial transplantation: Ratio of bone marrow chimerism in HSCs displayed as chimerism in secondary transplantation divided by chimerism in primary transplantation. Mice were reconstituted with 50% CD45.2^+^ CTR or cKO bone marrow cells mixed with 50% CD45.1^+^ wild-type bone marrow cells. (E-G) Long-term self-renewal and differentiation potential of 30 phenotypic HSCs analyzed during primary and secondary transplantation. (E) HSC chimerism; (F) CD19+ B cell chimerism; (G) Gr1+ granulocyte chimerism. Data in all panels are representative of two or more experiments. Bars and lines indicate average, circles depict individual mice. n.s., not significant; P > 0.05 by unpaired Student’s *t* test.

### Control of HSC numbers by differentiated hematopoietic cells is CXCR4 independent

The HSC compartment size is controlled by approximately 2-fold through feedback signals such as TGFβ, PF4, and adenosine, provided by differentiated hematopoietic cells (Bruns et al., 2014; Chow et al., 2011; Hirata et al., 2018; Zhao et al., 2014). *Cxcr4* deletion at the MPP stage ablates CXCR4 function in differentiated hematopoietic cells that control HSC numbers, namely megakaryocytes, macrophages, and Tregs (Bruns et al., 2014; Chow et al., 2011; Hirata et al., 2018; Zhao et al., 2014). To test if differentiated hematopoietic cells require CXCR4 for controlling the size of the HSC compartment, we conditionally deleted *Cxcr4* in differentiated immune cells downstream of the MPP stage (Fig. 3A) using multiple Cre-recombinase transgenic approaches (Abram et al., 2014; Schlenner et al., 2010; Tiedt et al., 2007). We found no measurable differences in HSC numbers, frequency, and cell cycle status in the bone marrow of mice in which *Cxcr4* was deleted in megakaryocytes (Fig. 3B and Fig. S2A), macrophages (Fig. 3C and Fig. S2B), lymphoid cells (Fig. 3D and Fig. S2C), and T cells (Fig. 3E and Fig. S2D). In mice conditionally deficient in CXCR4 in T cells, the number of bone marrow Tregs was significantly reduced and to a greater extent than in MPP CXCR4 cKO mice (Fig. S2E). In mice conditionally deficient in CXCR4 in megakaryocytes or in myeloid cells, the number of bone marrow megakaryocytes, neutrophils, monocytes, and CD160+ macrophages was equivalent to that in control littermate mice (Fig. S2F and G). Thus, the increased numbers of HSCs seen in mice in which MPPs lack *Cxcr4* is not due to the lack of CXCR4 expression in differentiated hematopoietic cells, including in Tregs (Hirata et al., 2018).

**Figure 3.**
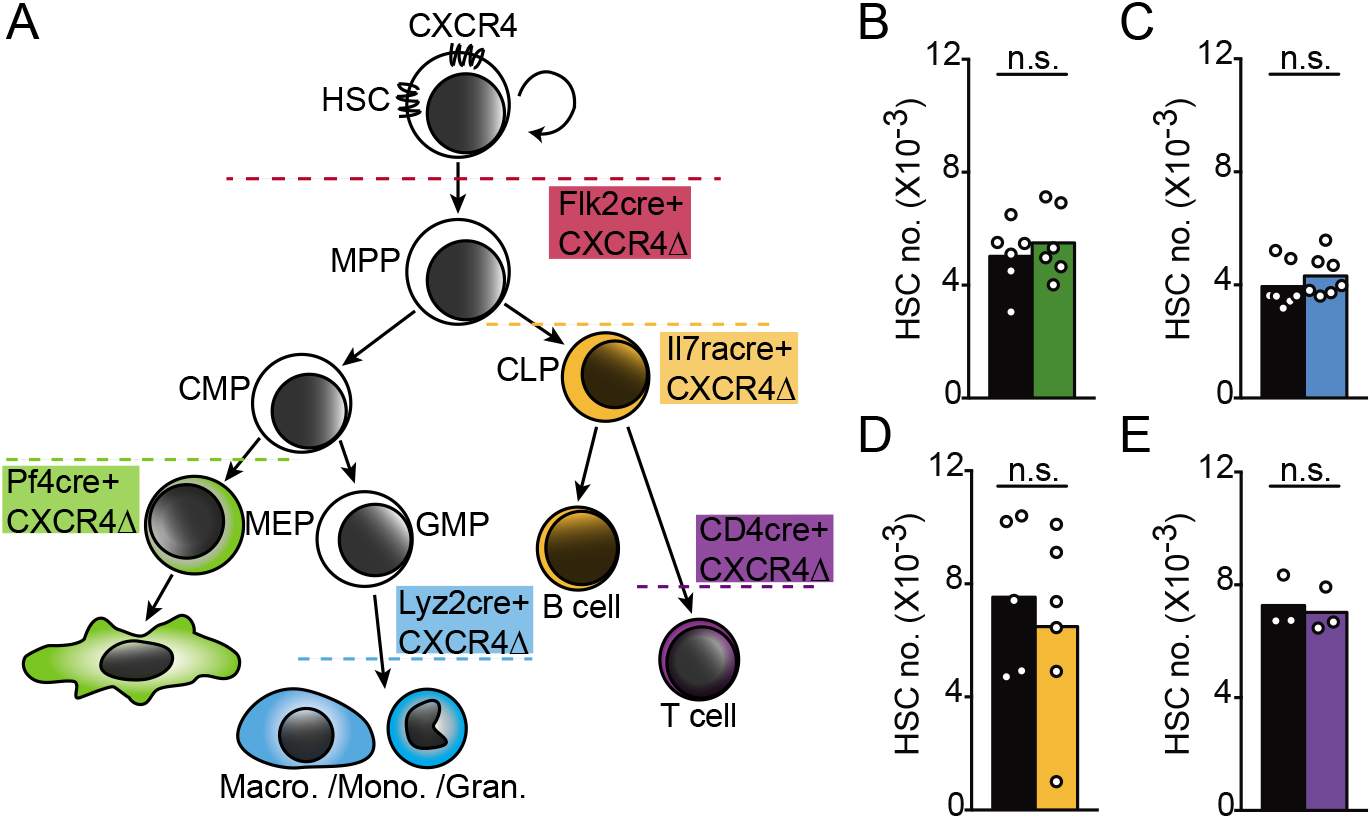
Normal HSC numbers in mice carrying *Cxcr4* deletion in megakaryocytes, myeloid, and lymphoid cells. (A) Schematic representation of CXCR4 deletion in hematopoietic cells by cell-lineage specific cre-recombinase transgenic approaches. (B-E) LT-HSC numbers in one femur and tibia of control littermate mice (black); (B) *Pf4-cre.Cxcr4^fl/fl^* (green); (C) *Lyz2-cre.Cxcr4^fl/fl^* (blue); (D) *Il7ra-cre.Cxcr4^fl/fl^* (yellow); (E) *Cd4-cre. Cxcr4^fl/fl^* (purple). Data are representative of two or more experiments. Bars indicate average, circles depict individual mice. n.s., not significant, P > 0.05 by unpaired Student’s *t* test.

### Stability of HSC niche cell transcriptional heterogeneity

MSPCs and ECs are critical regulators of HSC homeostasis and also provide a myriad of signals required for hematopoietic progenitor differentiation (Miao et al., 2020). The ability of MSPCs and ECs to produce hematopoietic cytokines and chemokines can be altered under pre-leukemic and leukemic states (Baryawno et al., 2019; Fistonich et al., 2018; Miao et al., 2020; Zehentmeier and Pereira, 2019). To determine if the HSC niche is altered when MPPs lack *Cxcr4*, we analyzed the transcriptional heterogeneity of non-hematopoietic bone marrow cells by droplet-based single-cell RNA sequencing (scRNA-seq). Total bone marrow cells were isolated from crushed femurs and tibias, and single cell suspensions were prepared after a brief 30-minute treatment with collagenase. Cells were sorted based on the negative expression of hematopoietic cell-specific membrane-associated proteins by fluorescence-activated cell sorting (Fig. S3A), and mRNA libraries (10x Genomics) were prepared and sequenced (see Methods). A total of 14,027 cells, of which 6,413 were from control and 7,614 from *Cxcr4* cKO mice, were profiled at a mean depth of 39,451 and 51,660 reads/cell, respectively. After quality control utilizing Seurat (Stuart and Butler et al., 2019) (see Methods) and removal of contaminating hematopoietic cells, we analyzed 1,380 control and 2,438 cKO cells for a total of 3,818 non-hematopoietic cells. We used the dimensional reduction technique Uniform Manifold Approximation and Projection (UMAP) to visualize non-hematopoietic cell clusters and compare differences in cell cluster heterogeneity between control and cKO bone marrow samples (McInnes, 2018). Unsupervised clustering identified 2 mesenchymal lineage cell clusters marked by *Lepr* expression, 3 osteolineage clusters, 4 endothelial cell clusters, 1 fibroblast cell cluster, and 1 chondrocytic cell cluster in both datasets (Fig. 4A and B, and Fig. S3B and C). The total number of stromal and endothelial cell clusters resolved was reduced when compared with those described in prior studies (Baccin et al., 2020; Baryawno et al., 2019; Tikhonova et al., 2019), possibly because of a reduced number of cells sequenced. Nevertheless, the overall structure of the data was comparable: one cycling EC cluster (cluster 11), two major cell clusters representing arterial and arteriolar ECs (clusters 10 and 2, respectively), and one sinusoidal EC subset (cluster 1); a large population of Lepr+ mesenchymal lineage cells (cluster 0); and a small cluster of Lepr^+^ mesenchymal lineage cells with increased expression of immediate early genes (e.g., *Fos, Fosb, Jun, Nr4a1, Mcl1*, cluster 7) (Tullai et al., 2007). Although cluster 9 is transcriptionally similar to cluster 1, cluster 9 cells express very low amounts of EC-specific genes (Fig. 4B shows *Cdh5*, but similar results could be seen for *Kdr, Flt4*, and *Flt1* expression). Comparison of cell clusters identified in control and cKO bone marrow samples revealed considerable overlap between samples (Fig. 4C), and a comparable proportion of individual clusters (Fig. 4D). Differential abundance test between control and cKO datasets using Milo (Dann et al., 2020) confirmed that the abundance of cellular states is not substantially different in all clusters (Fig. 4E; spatial False Discovery Rate (FDR) < 0.05). Furthermore, we found a very small number of differentially expressed genes (DEGs) between control and cKO mesenchymal and endothelial cell clusters. The major Lepr^+^ cell population (cluster 0) with an adipocytic gene expression program previously identified (Baccin et al., 2020) showed 11 DEGs with LogFC < 1 (Table 1). Likewise, the major sinusoidal and arteriolar EC clusters revealed 23 and 11 DEGs with LogFC < 1, respectively (Table 1). Mesenchymal and osteolineage cells (clusters 3-7, Fig. S3D) also revealed < 15 DEGs in which differences between samples did not exceed two-fold (Table 1). Importantly, none of the DEGs have been implicated in HSC regulation nor in downstream hematopoietic progenitor differentiation, and no major biological pathway could be associated with DEGs revealed in these analyses. Major regulators of HSC homeostasis such as *Cxcl12, Kitl, Ptn*, and adhesion ligands *Vcam1* and *Icam1* were transcriptionally abundant and equivalent in mesenchymal and endothelial cell clusters of both samples (Fig. 4F). Furthermore, no differences were detected in hematopoietic growth factors expressed in both samples (Fig. S3F). Finally, to test if the bone marrow environment of cKO mice can support HSCs we transplanted WT BM into lethally irradiated WT or cKO recipient mice. We found similar numbers of phenotypic HSCs in the bone marrow of WT and cKO recipient mice 16 weeks after transplantation (Fig. S3G). Taken together, these studies suggested strongly that major transcriptional changes in HSC niches could not explain the HSC expansion seen in cKO mice, and that the cKO bone marrow environment *per se* was insufficient for causing HSC expansion.

**Figure 4.**
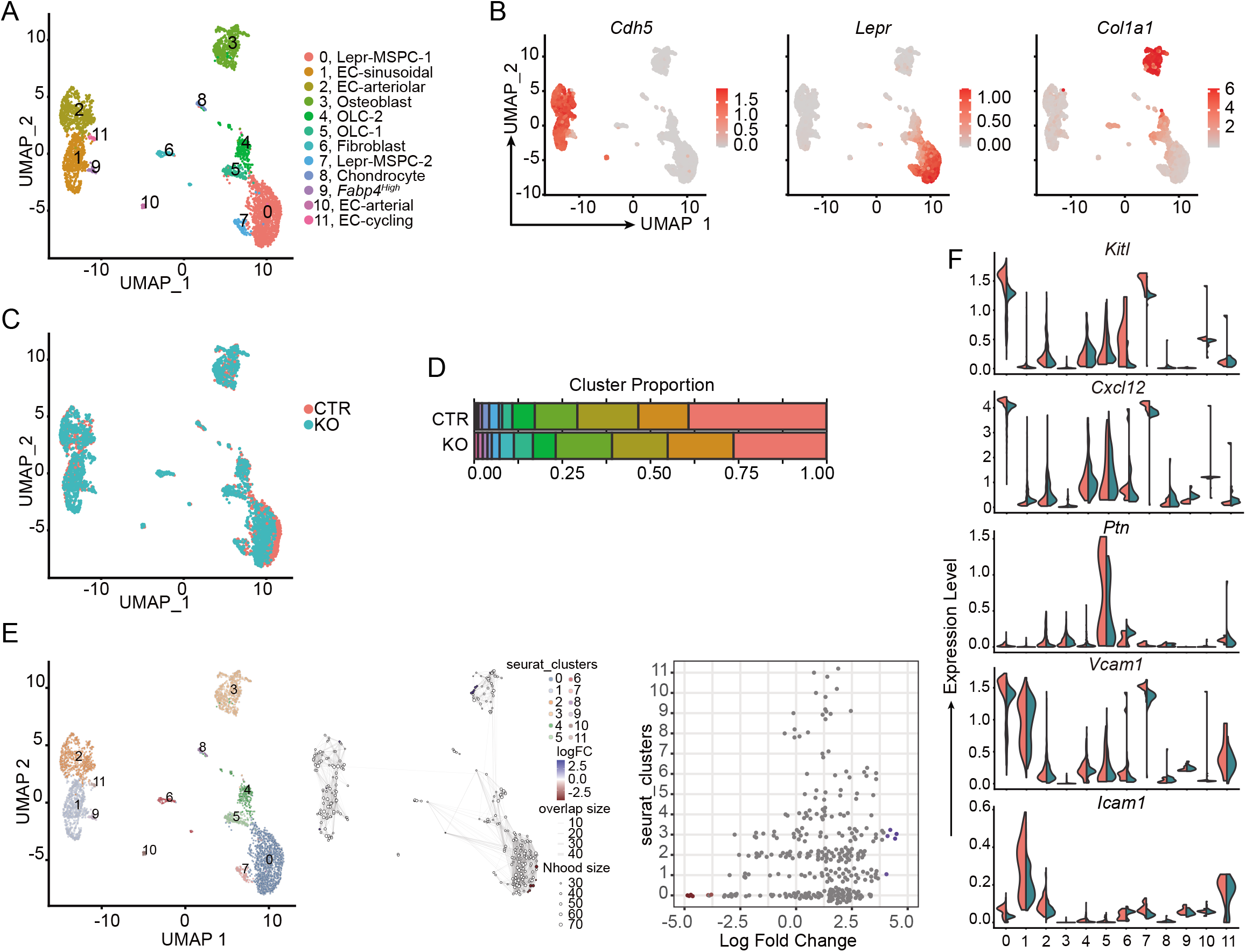
Niche cell transcriptional heterogeneity in mice with CXCR4-deficient MPPs. (A) UMAP visualization of bone marrow non-hematopoietic cells. (B) Expression levels of genes associated with endothelial cells (*Cdh5*), MSPCs (*Lepr*), and osteolineage cells (*Col1a1*) overlaid on UMAP. (C) Overlay of UMAP visualization of bone marrow non-hematopoietic cells from *Flk2-cre.Cxcr4^fl/+^* (CTR, red) and *Flk2-cre.Cxcr4^fl/fl^* (KO, blue) mice. (D) Cluster proportion. (E) Differential abundance test with Milo. UMAP cluster representation (left); graph representation of Milo differential abundance testing (middle); Beeswarm plot showing the distribution of logFC in neighborhoods containing cells from different Seurat clusters (right). Nodes are neighborhoods colored by logFC between CTR and KO. Non-differential abundance neighborhoods (FDR 5%) are colored white; neighborhood sizes correspond to number of cells in each neighborhood. Graph edges represent the number of cells shared between adjacent neighborhoods. Cell cluster frequencies in each sample (horizontal-colored bars). (F) Violin plots representing expression levels of essential HSC regulators (*Kitl, Cxcl12, Ptn, Vcam1, Icam1*) in CTR (red) and KO (blue) cell clusters.

To increase the sensitivity of detecting DEGs, we performed RNA sequencing of bulk sorted PDGFRα^+^ MSPCs, which greatly overlap with LEPR^+^ MSPCs (not shown). This analysis revealed 53 DEGs (normalized RNA counts >0 in all samples, q-value<0.05) (Fig. 5A-C), of which none are hematopoietic cytokines, chemokines, nor genes known to directly or indirectly regulate HSC homeostasis (Fig. 5D). Thus, the bulk and scRNAseq studies revealed a remarkable stability of niche cell transcriptome and niche cell heterogeneity under conditions in which hematopoietic progenitors and differentiated cells interact poorly with CXCL12-producing niche cells.

**Figure 5.**
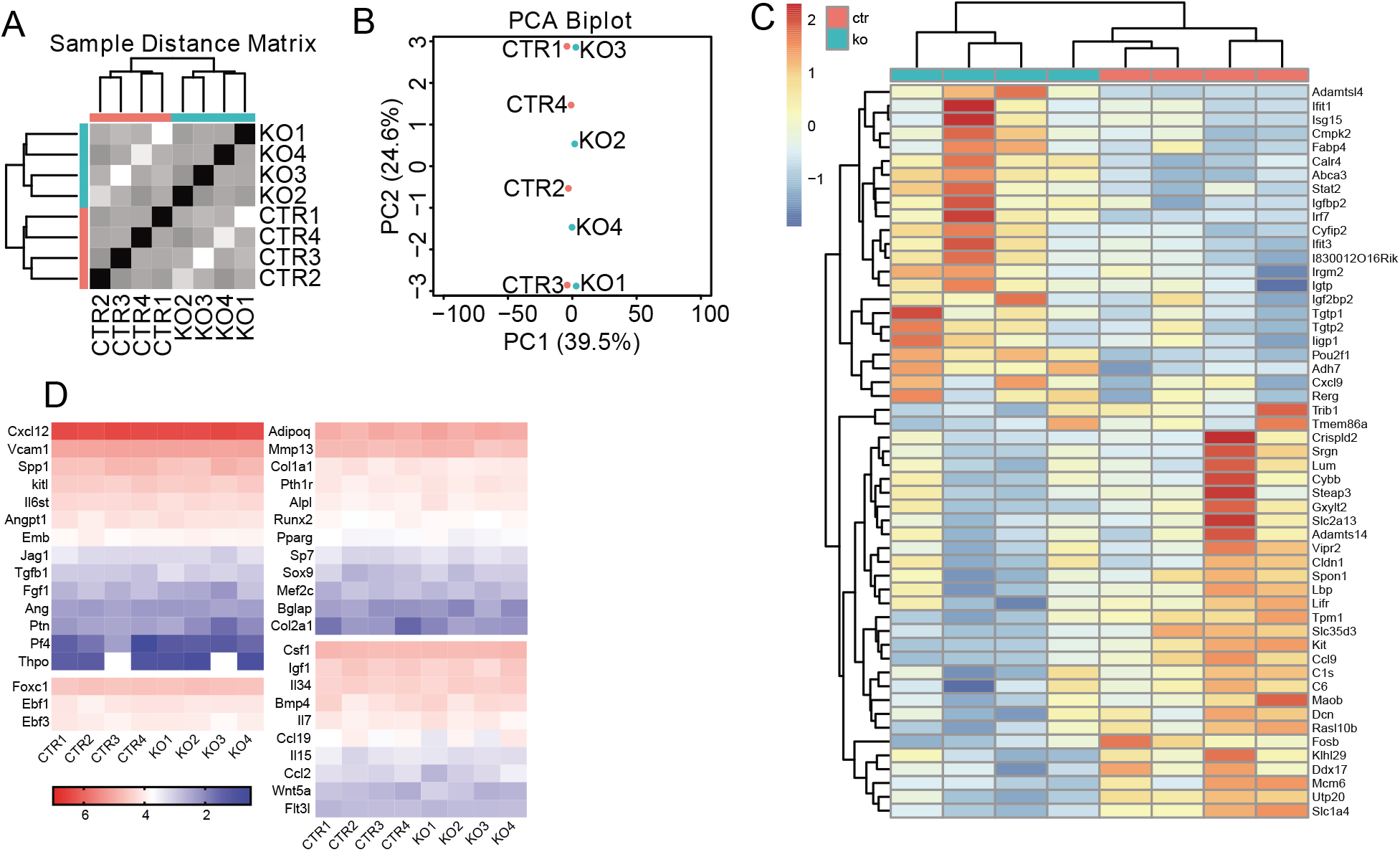
Comparison of MSPC global transcriptome between control and mice with CXCR4-deficient MPPs. (A) Unsupervised hierarchical sample clustering represented as Sample Distance Matrix. CTR (*Flk2-cre.Cxcr4^fl/+^*) and KO (*Flk2-cre.Cxcr4^fl/fl^*). (B) Principal-component analysis (PCA) representation of *Flk2-cre.Cxcr4^fl/+^* (CTR) and *Flk2-cre.Cxcr4^fl/fl^* (KO) samples obtained from individual mice. (C) Differentially expressed genes in MSPCs isolated from *Flk2-cre.Cxcr4^fl/+^* (red) and *Flk2-cre.Cxcr4^fl/fl^* (blue) mice. Plotted genes showed normalized RNA counts >0 in all samples, FDR adjust p<0.05. (D) Heatmap representation of log10 normalized RNA counts of genes important for HSC maintenance and multilineage differentiation.

### CXCR4 in hematopoietic progenitors controls HSC numbers through competition for membrane-bound SCF

Recent studies revealed discrepancy between HSC niche cell transcriptome and proteome (Asada et al., 2017; Severe et al., 2019). Thus, we considered the possibility that the abundance of hematopoietic growth factors displayed on WT and cKO niche cells may differ. Of note, SCF is particularly attractive given its critical role in HSC homeostasis, survival, and proliferation, and the fact that mSCF is particularly important for hematopoiesis (Anderson et al., 1990; Barker, 1994; Barker, 1997). Although transcript levels for SCF (*Kitl*) did not reveal differences between cKO and control Lepr+ MSPCs (Fig. 6A), we detected a significant increase in mSCF on bone marrow LEPR^+^ MSPCs and ECs (Fig. 6B-D) when staining with a rat anti-mouse SCF antibody (see Methods). We performed a series of in vitro and in vivo experiments to validate the specificity of mSCF staining. *Kitl* overexpression in OP-9 stromal cells showed robust mSCF expression relative to control transduced OP9 cells (Fig. S4A). Preincubation of the anti-SCF antibody with varying amounts of soluble SCF completely prevented mSCF detection on cKO MSPCs and ECs in a SCF dose-dependent manner (Fig. S4B). Importantly, reducing mSCF protein abundance by half through genetic means (i.e., crossing cKO mice with *Kitl^GFP/+^* mice), not only reduced mSCF protein levels in niche cells of cKO mice to the level detected in niche cells isolated from WT mice (Fig. 6E), but also reduced the number of phenotypic HSCs to physiological levels (Fig. 6F). Conditional *Kitl* deletion from LEPR^+^ MSPCs of *Cxcr4* cKO mice also reduced mSCF levels and HSC numbers (Fig. S4D and S6E, respectively). The levels of mSCF on MSPCs and ECs of mice conditionally deficient in *Cxcr4* in T cells (*Cd4*-Cre), myeloid cells (*Lyz2*-Cre), or in megakaryocyte (*Pf4*-cre) cells were normal (Fig. S4F and S4G). The total numbers of LEPR^+^ MSPCs and ECs were also equivalent between cell-lineage specific *Cxcr4* cKO mice and littermate controls (Fig. S4H and S4I). Besides SCF, HSCs also require hepatocyte-produced Thrombopoietin (Decker et al., 2018). Interestingly, we measured a small but significant reduction in Thrombopoietin concentration in the serum of cKO mice (Fig. S5A). However, heterozygous mutations in *Thpo* do not change the total number of HSCs in vivo, suggesting that long-range acting Thrombopoietin is not limiting the size of the HSC compartment (Decker et al., 2018).

**Figure 6.**
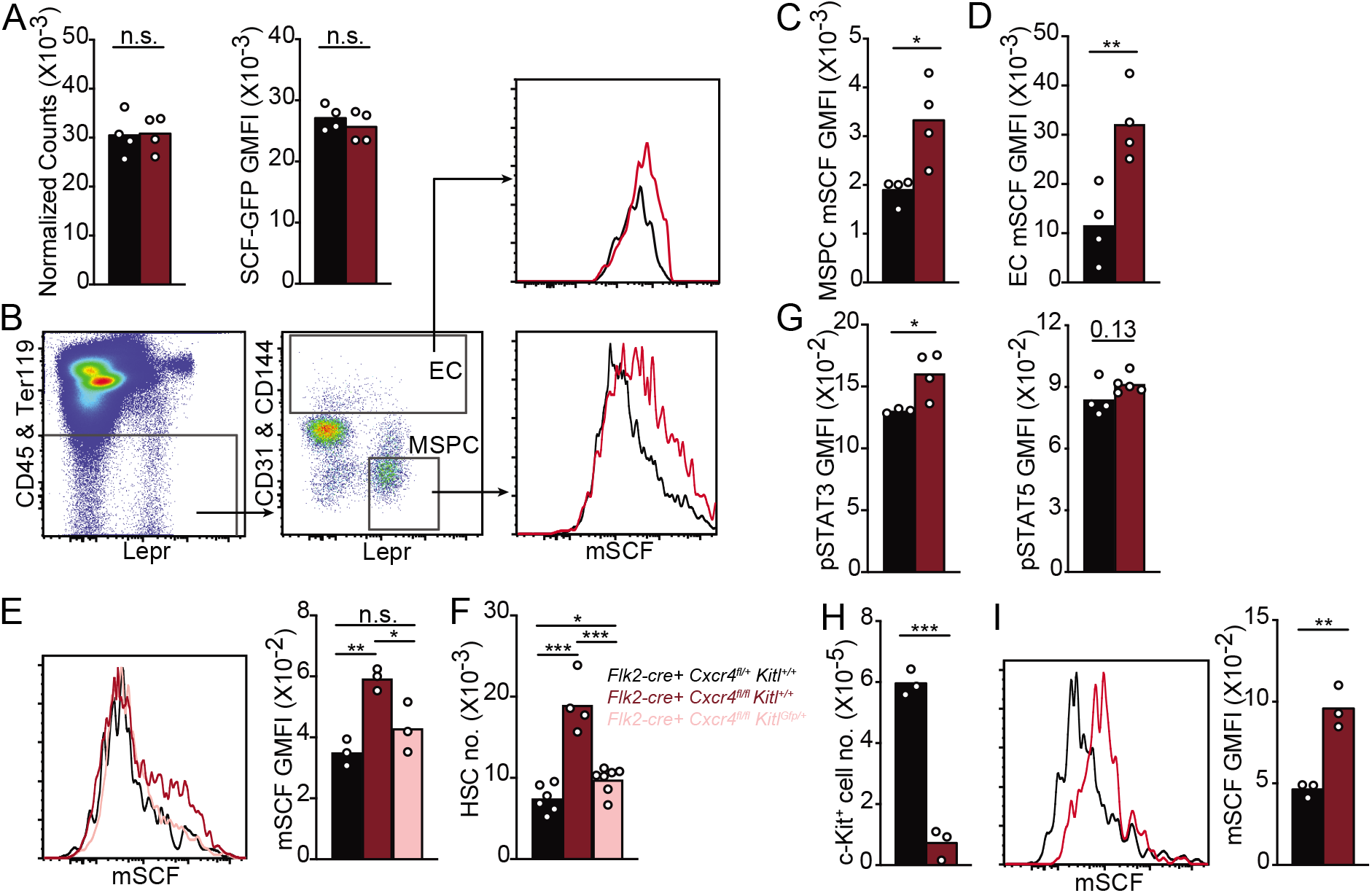
Altered membrane-bound SCF on MSPCs and ECs, and altered cKit signaling in HSCs of mice with CXCR4-deficient MPPs. (A) *Kitl* mRNA expression. Normalized *Kitl* RNA counts in MSPCs (left); and *Kitl-GFP* expression in gated LEPR^+^ MSPCs (right) of *Flk2-cre.Cxcr4^fl/+^ Kitl^GFP/+^* (black), and *Flk2-cre.Cxcr4^fl/fl^ Kitl^GFP/+^* (red) mice. (B) Gating strategy of LEPR^+^ MSPCs and ECs, and mSCF measurements by flow cytometry. (C and D) mSCF geometric mean fluorescent intensity (GMFI) on LEPR^+^ MSPCs (C) and ECs (D) from *Flk2-cre.Cxcr4^fl/+^* (black) and *Flk2-cre.Cxcr4^fl/fl^* (red) mice. (E) Overlay of mSCF on gated LEPR^+^ MSPCs isolated from indicated mice (left), mSCF quantification (right). (F) HSC numbers in bone marrow. Black, *Flk2-cre.Cxcr4^fl/+^Kitr^+/+^*; Red, *Flk2-cre.Cxcr4^fl/fl^ Kitl^+/+^;* Pink, *Flk2-cre.Cxcr4^fl/fl^ Kitl^GFP/+.^* (G) pSTAT3 (left) and pSTAT5 (right) levels in HSCs isolated from *Flk2-cre.Cxcr4^fl/+^* (black) and *Flk2-cre.Cxcr4^fl/fl^* (red) mice. (H) cKit^+^ cell number in bone marrow (I) mSCF levels on LEPR^+^ MSPCs. (H and I) Mice were treated with 200 μg of isotype (black) or ACK2 antibody (red) for 3 days. Data in panel A are from one bulk RNA sequencing experiment with n=4 samples/group. Data in panels B-I are representative of two or more experiments. Bars indicate average, circles depict individual mice. *, P < 0.05; **, P < 0.01 and *** P < 0.001 by unpaired Student’s *t* test.

cKit signaling activates the JAK/STAT pathway and induces STAT3 and STAT5 phosphorylation (Chung et al., 2006; Linnekin, 1999). HSCs stimulated with soluble SCF in vitro for 30 minutes revealed a low but significant increase in pSTAT3 and pSTAT5 measured by intracellular phospho-flow cytometry (Fig. S5C). Importantly, we could measure significantly increased pSTAT3 and a trend towards increased pSTAT5 in HSCs isolated from the bone marrow of *Cxcr4* cKO mice relative to that detected in control littermate (Fig. 6G). Combined, these data demonstrate that the increased HSC number seen in cKOs is dependent on increased mSCF availability on LEPR^+^ MSPCs and ECs in the bone marrow.

Finally, we aimed at understanding the mechanism underlying increased mSCF levels in HSC niche cells of cKO mice. Measurements of soluble SCF showed that cKO and WT mice have similar SCF concentrations in the bone marrow interstitial fluid (Fig. S5B), suggesting that the increased mSCF level detected is not due to reduced proteolytic cleavage from niche cells. Therefore, we considered the possibility that competition between HSCs and cKit^+^ hematopoietic progenitors might limit mSCF abundance in niche cells such that it influences the HSC compartment size. To test this possibility, we depleted cKit^+^ cells in vivo by administering saturating amounts of an anti-cKit antibody (clone ACK2, 200 μg/mouse injected i.v., Fig. S5D) that elicits antibody-dependent cellular phagocytosis (Chhabra et al., 2016), and measured changes in mSCF displayed on MSPCs and ECs in the bone marrow. Treated animals showed a ~ 6-8 fold reduction in cKit+ cells 3 days after the treatment (Fig. 6H). Remarkably, mSCF displayed on the surface of bone marrow MSPCs increased promptly in ACK2-treated animals relative to that detected in mice treated with isotype control antibody (Fig. 6I). These data demonstrate an inverse relationship between mSCF abundance on niche cells and the number of cKit^+^ hematopoietic stem and progenitor cells in the bone marrow. Combined, these data suggest strongly that mSCF acts as a carrying capacity factor that limits the size of the HSC compartment.

## Discussion

Guided by CXCR4, HSCs, hematopoietic progenitors, and differentiated immune cells localize in the bone marrow and physically interact with a heterogeneous population of cytokine and growth factor-producing mesenchymal and endothelial cells. While the lymphoid, myeloid, erythroid, and megakaryocyte lineages diverge in cytokine requirements at late stages of differentiation, they converge in their requirement for SCF during early developmental stages (Miao et al., 2020). Studies over the last 10 years identified LEPR^+^ mesenchymal lineage cells as relevant cellular sources of SCF for HSCs and lineage-restricted hematopoietic progenitors, including lymphoid, erythroid, and myeloid progenitors (Comazzetto et al., 2019; Ding et al., 2012). The LEPR^+^ MSPC population is transcriptionally and functionally heterogeneous and includes a major cell population characterized by an adipogenic gene expression program and smaller subsets of osteolineage-primed mesenchymal progenitors. Importantly, SCF produced by osteolineage-primed mesenchymal progenitors (marked by Osteolectin expression) contributes to CLP and early B and T cell progenitor development or maintenance, but it does not contribute to HSC homeostasis and early myeloid, erythroid, and megakaryocyte progenitor development in vivo (Shen et al., 2021). Therefore, HSCs and multiple hematopoietic progenitors rely on SCF produced presumably by the same fraction of MSPC that is marked by high *Lepr, Kitl*, and *Cxcl12* expression, a model that is in agreement with earlier findings that HSCs and MPPs are occasionally found in close proximity to the same niche cell (Cordeiro Gomes et al., 2016; Lim et al., 2017).

In this study, we uncovered an unexpected layer of regulation of HSC homeostasis. By displacing MPPs and downstream hematopoietic progenitors from CXCL12-producing niches in the bone marrow through conditional *Cxcr4* deletion, we made two unexpected observations. First, the HSC niche is transcriptionally stable and impervious to non-malignant cell-cell interactions. This finding contrasts with profound transcriptional changes that occur when niche cells interact with preB cell acute lymphoblastic leukemia or with acute myeloid leukemias, which impact hematopoiesis and MSPC differentiation in vivo (Baryawno et al., 2019; Fistonich et al., 2018; Mendez-Ferrer et al., 2020; Zehentmeier and Pereira, 2019). These observations suggest that signals provided or triggered by leukemic cells are sensed by HSC niche cells and that such signals control hematopoietic cytokine and chemokine production. Second, HSCs and SCF-dependent cKit-expressing hematopoietic progenitors compete for limited amounts of membrane-tethered SCF in vivo. This model of limited SCF availability is supported by the fact that *Kitl* haploinsufficiency reduces the number of HSCs and hematopoietic progenitors by approximately 2-fold under homeostatic conditions (Comazzetto et al., 2019; Ding et al., 2012).

Even though *Kitl* mRNA is abundantly expressed in MSPCs, we were unable to reliably detect mSCF protein on LEPR^+^ MSPCs of wild type mice when compared to the background staining detected in LEPR^+^ MSPCs of *Kitl* conditionally deficient mice (*Lepr-cre+ Kitl^fl/fl^*; Fig. S6C). However, mSCF protein was readily detected on MSPCs of wild type mice treated with a cKit blocking antibody (Fig. 6I). Furthermore, mSCF was also readily detected in MSPCs and ECs of mice in which hematopoietic progenitors interact poorly with niche cells (*Cxcr4* cKO mice; Fig. 6E and S4D). Combined, these studies provide evidence supporting a model in which competition for limited amounts of mSCF establishes a fine balance between the HSC and cKit^+^ hematopoietic progenitor compartment size. This model is in agreement with earlier studies by Nagasawa and colleagues showing that when large numbers of transplanted HSCs engraft into non-irradiated recipients, the number of host and donor-derived hematopoietic progenitors, such as GMPs, is reduced (Shimoto et al., 2017). It is also consistent with recent findings showing discrepancies between LEPR^+^ MSPC *Kitl* transcript and SCF protein abundance (Severe et al., 2019), while providing an alternative explanation: cytokine consumption by hematopoietic progenitors may account for such discrepancies.

It is presently unclear why HSCs do not expand by more than 2-fold when mSCF levels are still elevated in *Cxcr4* cKO mice. It is interesting to note that prior studies examining the HSC niche and mechanisms controlling the HSC compartment size also reported ~2-fold changes. For example, Angiogenin and Embigin have been shown to limit HSCs numbers by ~ 2-fold (Silberstein et al., 2016). Likewise, CXCL4 and TGFβ1 produced by megakaryocytes also limit HSC numbers by a similar magnitude. It is possible that the capacity for mSCF availability to modulate the HSC compartment size is limited by cues promoting HSC quiescence such as Angiogenin, Embigin, CXCL4 or TGFβ1. Future studies will determine if (and which) quiescence-promoting cues limit HSC growth under conditions of elevated mSCF availability.

The exact mechanism by which HSCs and hematopoietic progenitors consume mSCF remains unclear. Recent studies using intravital 2-photon microscopy revealed that HSCs are moderately dynamic under homeostasis (Christodoulou et al., 2020; Upadhaya et al., 2020). Like developing B cells, HSC motility is dependent on active CXCR4 signaling, and HSC retention within bone marrow requires integrin-mediated adhesion (Beck et al., 2014; Upadhaya et al., 2020). Likewise, MPPs are also motile within bone marrow under transplantation, and also require CXCR4 for bone marrow retention (Cordeiro Gomes et al., 2016; Lo Celso et al., 2009). It is possible that interactions between HSCs, hematopoietic progenitors, and SCF-producing niche cells result in the physical removal of mSCF through a process resembling trogocytosis (Dance, 2019; Joly and Hudrisier, 2003). Future studies that allow visualization of mSCF on bone marrow niche cells may provide insights into the mechanism(s) that limit mSCF availability in vivo.

Cell competition for limiting resources is common and physiologically important. In the immune system, examples include B cell clonal competition during the germinal center reaction, competition for access to B lymphocyte survival cytokines such as BAFF, among several others (Cyster et al., 1994; Victora and Nussenzweig, 2012). In these cases, competition ensures the survival of the fittest cell or clone with direct benefit for the host (e.g., selection of high-affinity clones, elimination of autoreactive lymphocytes). Likewise, competition for limited amounts of mSCF may ensure that the fittest HSCs are maintained and contribute to blood cell production over time. Once HSCs divide and progressively differentiate into MPP and lineage-restricted progenitors, their shared dependency on mSCF may ensure homeostatic control of HSC and progenitor compartment sizes. These findings and model are more easily compatible with studies showing that HSCs actively contribute to blood cell production during homeostasis (Sawai et al., 2016) than with other studies suggesting that HSCs contribute only during physiological stress conditions such as systemic infection or inflammation (Busch et al., 2015).

A recent study demonstrated that Tregs require CXCR4 for homing to the bone marrow where they play a direct role in controlling HSC homeostasis by secreting adenosine (Hirata et al., 2018). However, we failed to measure significant changes in the HSC compartment size when T cells lack CXCR4, including Tregs. A major difference between these studies is that Hirata et al. analyzed mice in which only Tregs are *Cxcr4*-deficient whereas in our studies we deleted *Cxcr4* in all T cell subsets. Under homeostasis, naïve T cells express very little CXCR4 and are largely unable to migrate into the bone marrow (Arojo et al., 2018). In contrast, memory T cells upregulate CXCR4 and migrate into the bone marrow where these cells receive homeostatic signals (presumably IL7 and/or IL15) required for their long-term maintenance (Chaix et al., 2014; Tokoyoda et al., 2009). Thus, it is possible that adenosine produced by bone marrow Tregs suppresses bystander activation of recirculating memory T cells in order to reduce or prevent the release of inflammatory signals in the HSC niche (Kim and Shin, 2019).

In summary, our studies revealed that mSCF acts as a carrying capacity factor for HSC homeostasis due to competition from cKit-dependent hematopoietic progenitors for limited amounts of mSCF under homeostatic conditions. Furthermore, while MSPCs respond to long-range cues such as hormones for regulation and differentiation, these cells and ECs are impervious to alterations in cell-cell interactions with hematopoietic progenitors and remain transcriptionally stable.

## Data Availability Statement

All bulk RNA-seq and scRNA-seq data are being deposited in the Gene Expression Omnibus (GEO) under the accession number GSE171015.

## Acknowledgements

We thank Vivian Y. Lim, Abigail Jarret, and Tianyang Mao (Yale University) for help with some experiments; Aaron Ring and Shangqin Guo (Yale University) for insightful suggestions and technical support; and Sean J. Morrison and Stefano Comazzetto (UT Southwestern) for sharing biological samples. These studies were funded by the NIH (RO1AI113040 and R21AI13306001A1 to J.P. Pereira), and by the National Research Foundation of Korea (NRF) grant funded by the Korea government (MSIT) (No. 2020R1F1A1076705).

## Methods Details

### Mice

Adult C57BL/6NCR (strain code 556) (CD45.2+) and B6-Ly5.1/Cr (stain code 564) (CD45.1+) were purchased from Charles River Laboratories. *Pf4*-cre, *Kitl^fl/fl^, Lepr*-cre, and *Kitl^GFP/+^* mice were purchased from The Jackson Laboratories. *Cxcr4^fl/fl^, Lyz2*-cre, *Il7ra*-cre, *CD4*-cre, and *Rosa26t^dtomato/+^* mice were from internal colonies. *Flk2*-cre mice were a gift from Dr. E. Camilla Forsberg (University of California, Santa Cruz). Although *Flk2*-cre transgene is inserted into Y-chromosome, our previous work showed the hematopoietic cell composition in bone marrow and secondary lymphoid organs of *Flk2*-cre transgenic mice is indistinguishable from their littermate controls. All mice were maintained under specific pathogen-free conditions at Yale Animal Resources Center and were used according to the protocol approved by the Yale University Institutional Animal Care and Use Committee.

### Flow cytometry

For analyses of hematopoietic cell composition, bone marrow cells were obtained by crushing long bones with DMEM supplemented with 2% FBS, 1% Penicillin/Streptomycin, 1% L-glutamine, and 1% HEPES. Spleen cells were obtained by mashing spleens through 70 μm cell strainers with the same media. Bone marrow stromal cell isolation and mSCF staining were performed as previously described (Cordeiro Gomes et al., 2016). Cells were counted with a Beckman Coulter Counter. Cells were then stained with antibody cocktail diluted in FACS buffer, at the concentration of 25μl per 1 × 10^6^ cells on ice. To stain intracellular proteins including Ki67, pSTAT3, and pSTAT5, cells were fixed in Cytofix/Cytoperm solution (BD) for 20 min at a concentration of 25μl per 1 × 10^6^ cells on ice and washed twice with PermWash buffer (BD). Cells were then permeabilized in Permeabilization Buffer Plus (BD) at the same concentration on ice for 10 min and washed twice. The Ki67 antibody was diluted in FACS buffer and was used to stain cells on ice. The pSTAT3 and pSTAT5 antibodies were diluted in PermWash buffer, and the cells were stained at room temperature. Hematopoietic cell populations were identified as follows: LSK: Lineage^-^ cKit^+^ SCA-1^+^; LT-HSC: Lineage^-^ cKit^+^ SCA-1^+^ FLT3^-^ CD150 (SLAM)^+^; ST-HSC: Lineage^-^ cKit^+^ SCA-1^+^ FLT3^-^ CD150^-^; MPP: Lineage^-^ cKit^+^ SCA-1^+^ FLT3^+^ CD150^-^ (The lineage cocktail was: CD19, B220, CD3e, CD4, Gr1, NK1.1, Ter119, CD11b, CD11c, CD41, CD48). MPP2: Lineage^-^ cKit^+^ SCA-1^+^ FLT3^-^ CD48^+^ SLAM^+^; MPP3: Lineage^-^ cKit^+^ SCA-1^+^ FLT3^-^ CD48^+^ SLAM^-^; MPP4: Lineage^-^ cKit^+^ SCA-1^+^ FLT3^+^ (The lineage cocktail was: CD19, B220, CD3e, CD4, Gr1, NK1.1, Ter119, CD11b, CD11c). Treg: CD3e^+^ CD4^+^ CD25^+^; MSPC: CD45^-^ Ter119^-^ CD31^-^ CD144^-^ LEPR^+^; EC: CD45^-^ Ter119^-^ CD31^high^ CD144^high^.

### Immunostaining and Microscopy analyses

Freshly dissected femurs were fixed in 2% paraformaldehyde-based fixative in PBS at 4°C overnight. Bones were dehydrated in a solution of 30% sucrose in PBS, at 4°C overnight. Samples were embedded in OCT and snap-frozen in an ethanol/dry ice bath. Frozen sections were prepared according to the Kawamoto method or using the CryoJane tape transfer system (Leica). Femur whole mounts frozen in OCT were stained with primary antibodies for 2–3 days at 4°C and secondary antibodies for 1 day at 4°C. Slides were mounted with Fluormount-G (SourthernBiotech) or with a 30% glycerin solution. Images were acquired on a Leica SP8 confocal microscope.

### Competitive reconstitution assay

Recipient mice (CD45.1^+^) were exposed to whole-body lethal irradiation from a ^137^Cs source (two doses of 550 rads separated by 3 h). Mice were then reconstituted with 5 × 10^6^ donor whole bone marrow cells (a mixture of CD45.2^+^ and CD45.1^+^ cells in a 1:1 ratio) by intravenous injection. Chimeric mice were analyzed 16 weeks after reconstitution. For long-term reconstitution analysis, 6 × 10^6^ donor whole bone marrow cells were injected to recipients, and the chimerism was analyzed 16 weeks after reconstitution in both primary and secondary transplantations.

### Bone marrow stromal cells preparation for bulk RNA sequencing

Long bones were flushed with HBSS supplemented with 2% of FBS, 1% Penicillin/Streptomycin, 1% L-glutamine, 1% HEPES, and 200 U/mL Collagenase IV (Worthington Biochemical Corporation). Cells were first digested at 37°C for 20 min and gently pipetted to dissociate cell clumps. Cells were then filtered through 100 μm cell strainers after being digested at 37°C for another 10 min, and washed with media. Stromal cells were enriched by depleting hematopoietic cells with biotin-conjugated CD45 and Ter119 antibodies, and Dynabeads® Biotin Binder (Invitrogen #11047). Cells were stained with CD31, CD144 and PDGFRα antibodies after depletion and CD45^-^ Ter119^-^ CD31^-^ CD144^-^ PDGFRα^+^ SCF-GFP^+^ cells were sorted into DMEM with 10% FBS. Sorting was performed by the BD FACS Aria II. The sorted cells were then sorted with the same gating strategy again to reduce contamination into 350 μL RLT plus buffer with 3.5 μL β-mercaptoethanol. RNA was extracted using the RNeasy® Plus Micro Kit (Qiagen #74034).

### Bulk RNA sequencing

RNA sequencing was performed using the Illumina HiSeq2500 system with paired-end 2 x 76 bp read length by the Yale Center for Genome Analysis. The sequencing reads were trimmed by 7 bp on the 5’-end and until QS≥20 on the 3’-end. The sequencing reads were then aligned onto the reference genome Mus musculus GRCm38 (mm10) using HISAT2 (Pertea et al., 2016) and converted to BAM files using SAMtools (Li et al., 2009). Read counts were generated using HTSeq-count (Anders et al., 2015) with GENCODE v27 as the gene model. By utilizing DESeq2 (Love et al., 2014), the read counts were normalized by size factor and the differential expression of genes were calculated using LRT model in correction for batch effect.

### Bone marrow stromal cells preparation for single-cell RNA sequencing

Bone marrow stromal cells were isolated using the same method as the bulk RNA sequencing cell preparation. The bones after flushing were chopped into small pieces and digested with HBSS supplemented with 2% of FBS, 1% Penicillin/Streptomycin, 1% L-glutamine, 1% HEPES, and 200 U/mL Collagenase IV (Worthington Biochemical Corporation) at 37°C for 45 min under agitation (120rpm). Cells were then filtered through 100 μm cell strainers before combined with digested bone marrow stromal cells. Cells were then stained with CD31, CD144, lineage (B220, CD19, CD11b, Gr1, CD3e) and CD71 antibodies. CD45^-^ Ter119^-^ Lin^-^ CD31^-^ CD144^-^ CD71^-^ cells were sorted by the BD FACS Aria II machine into 350 μL DMEM with 20% FBS. Single-cell RNA sequencing was performed by the Yale Center for Genome Analysis. The libraries were prepared using the Chromium Single Cell 3□ Reagent Kits v3 according to the protocol. Libraries were run on an Illumina NovaSeq system with 100-bp paired-end reads to ~80% saturation level and to get the coverage to ~ 40,000 reads per cell, and the sequencing reads were aligned onto Mus musculus GRCm38 (mm10) reference genome.

### scRNA-seq data preprocessing and analysis

Barcode processing and single cell 3’ gene count matrix calculation were conducted using the 10x Genomic Cell Ranger 4.0.0 (Zheng et al., 2017) and Gene-Barcode matrix containing 14,027 cells were generated. Quality control, finding highly variable genes, dimensionality reduction, graph-based unsupervised clustering, and identification of differentially expressed genes were performed using Seurat R Package 4.0 (Stuart et al., 2019). Cells having fewer than 200 or more than 6,000 detected features and more than 20% mitochondrial gene mapped reads were excluded from downstream analyses. Also, mitochondrial and ribosomal protein features were removed from the count matrix and contaminating hematopoietic cells expressing the genes listed in Table 2 were filtered out. Then the count data was normalized, and variance stabilized using the SCTransform (Hafemeister and Satija, 2019).

### Dimensionality reduction, unsupervised cell clustering, and visualization

With the normalized gene-barcode matrix of resulting 3,818 cells, highly variable genes were detected and dimensionality reduction with principal component analysis (PCA) was conducted. The number of optimal principal components (PCs) was determined by *ElbowPlot* function of Seurat. Dimensionality was reduced and projected the cells in 2D space using UMAP by *RunUMAP* function of Seurat. Markov affinity-based graph imputation of cells (van Dijk et al., 2018) was used to denoising the high-dimensional scRNA-seq data by imputing plausible gene expression in each cell.

### Annotation of cell types and identification of cluster markers

The markers defining each cluster were identified by performing *FindAllMarkers* in the Seurat R package using the MAST method (Finak et al., 2015). Feature plots with the top 10 significant positive markers of each cluster and a heatmap of the top 10 significant positive markers were generated to visualize how well the clusters are defined. Also, DEGs between conditions within the same cluster were identified using *FindMarkers* with the MAST method.

### Differential Abundance test

Differential abundance of neighborhoods was analyzed via MiloR R package (https://github.com/MarioniLab/miloR) by allocating cells to partially overlapping neighborhoods on a k-nearest neighbor(KNN) graph. With calculated log2 fold change and FDR, clusters having statistically significant differential abundance were determined.

### Elisa Assays

To harvest bone marrow interstitial fluid, long bones were flushed into 1.5ml Eppendorf tubes using compressed air and immediately weighed and dissociated in 15-fold DPBS by pipetting and vortexing. The mixture was then centrifuged at 1200 rpm for 5 min, and the supernatant was transferred to new tubes, followed by another centrifugation at 10,000 rpm for 5 min to remove the remaining debris. To harvest serum, blood was collected and allowed to clot at room temperature for 1 hour. The clotted blood was then centrifuged at 1500 rpm for 10 min and the serum was transferred to new tubes. The serum was diluted 5-fold for the assay. ELISA for SCF and TPO was conducted with Mouse SCF Quantikine ELISA Kit (R&D, MCK00) and Mouse Thrombopoietin Quantikine ELISA Kit (R&D, MTP00).

---

**Figure S1. *Flk2*-cre is activated at the MPP stage.**

(A) Gating strategies for bone marrow cell populations. (B) Enumeration of LT-HSC, ST-HSC, MPP cell subsets, erythroid, myeloid, and lymphoid progenitors, B cell progenitors, and monocytes and neutrophils in femur and tibia of *Flk2-cre.Cxcr4^fl/+^* (black) and *Flk2-cre.Cxcr4^fl/fl^* (red) mice. (C) Chimerism of lineage positive cells and MPPs in bone marrow of lethally irradiated mice reconstituted with 50% CD45.2^+^ *Flk2-cre.Cxcr4^fl/+^* (black) or *Flk2-cre.Cxcr4^fl/fl^* (red) bone marrow cells mixed with 50% CD45.1^+^ wild-type bone marrow cells. Bars indicate average, circles depict individual mice. *, P < 0.05; **, P < 0.01, ***, P < 0.001, and ****, P < 0.0001 by unpaired Student’s *t* test.

**Figure S2. HSC frequency and quiescence in *Cxcr4* conditionally deficient mice.**

(A, B and D) HSC and MPP4 cell frequency; HSC cell cycle status Control littermates (black), *Pf4-cre.Cxcr4^fl/fl^* (green), *Lyz2-cre.Cxcr4^fl/fl^* (blue) and *Cd4-cre.Cxcr4^fl/fl^* (purple) mice. (C) HSCs and MPP4 cell frequency in bone marrow of control (black) and *Il7ra-cre.Cxcr4^fl/fl^* (yellow) mice. (E) T regulatory (Tregs) cell number per femur and tibia: WT, wild-type; *Flk2-cre.Cxcr4^fl/fl^* (red), *Cd4*-cre.*Cxcr4^fl/fl^*. (F) Megakaryocyte numbers in *Pf4-cre.Cxcr4^fl/+^* (black) and *Pf4-cre.Cxcr4^fl/fl^* (green) mice. Megakaryocytes were enumerated by fluorescence microscopy analysis of 7 μm thick femur sections (left panel). Scale bar is 50μm. (G) Myeloid cell numbers in bone marrow of *Flk2-cre.Cxcr4^fl/fl^* (red), *Lyz2-cre.Cxcr4^fl/+^* (black) and *Lys2-cre.Cxcr4^fl/fl^* (blue) mice. Bars indicate average, circles depict individual mice. n.s., not significant, P > 0.05; *, P < 0.05 and *** P < 0.001 by unpaired Student’s *t* test.

**Figure S3. Niche cell heterogeneity and gene expression analyses.**

(A) Gating strategy used for sorting of bone marrow non-hematopoietic cells. (B) Cluster signature genes. Expression (row-wide Z-score of ln(TP10K+1)) of top differentially expressed genes (rows) across the cells (columns) in each cluster (color bar, top, indicates each cluster as in Fig. 4A). Genes are indicated on the left. (C) Expression levels of indicated cluster-defining genes overlaid on UMAP. (D) Osteolectin expression level overlaid on UMAP. (E) Expression level of the indicated genes in cell clusters of CTL and KO with respect to their pseudotime coordinates. Black lines depict LOESS regression fit of the normalized expression values of essential HSC regulators (*Kitl, Cxcl12, Ptn, Vcam1, Icam1*)in different clusters. (F) Violin plots representing expression levels of hematopoietic cell differentiation genes in CTL (red) and KO (blue) cell clusters. In panels E and F, CTL (*Flk2-cre.Cxcr4^fl/+^*) and KO (*Flk2-cre.Cxcr4^fl/fl^*). (G) HSC numbers in *Flk2-cre.Cxcr4^fl/+^* (CTR, black) and *Flk2-cre.Cxcr4^fl/fl^* mice (cKO, red) mice lethally irradiated and reconstituted with wild-type bone marrow cells. n.s., not significant, Student’s *t* test.

**Figure S4. mSCF measurements and staining specificity.**

(A) Histogram overlay of mSCF staining in OP9 cells transduced with Empty-Vector pMSCV (gray) or with *Kitl*-expressing pMSCV (red) (B) histograms in gated LEPR^+^ MSPCs (left) and ECs (right) of secondary antibody (gray), or of anti-SCF pre-incubated for 30 minutes at room temperature with 0 nM (red), 0.105 nM (blue), or 0.421 nM (dark gray) mouse recombinant SCF. (C) mSCF histogram overlay of LEPR^+^ MSPCs isolated from *Lepr*-cre^+^ *Kitl^fl/+^* (black) and *Lepr*-cre^+^ *Kitl^fl/fl^* (pink) mice. (D) mSCF histogram overlay of LEPR^+^ MSPCs isolated from *Lepr*-cre^+^ *Kitl^fl/+^ Flk2-cre^+^ Cxcr4^fl/fl^* (black) and *Lepr*-cre^+^ *Kitl^fl/f^ Flk2-cre^+^ Cxcr4^fl/fl^* (orange) mice. (E) HSC numbers in bone marrow of *Lepr*-cre^+^ *Kitl^fl/+^ Flk2-cre^+^ Cxcr4^fl/fl^* (black) and *Lepr*-cre^+^ *Kitl^fl/fl^ Flk2-cre^+^ Cxcr4^fl/fl^* (orange) mice. (F and G) mSCF expression on LEPR^+^ MSPCs (F) and ECs (G); (H and I) LEPR^+^ MSPC and ECs cell numbers per femur and tibia. In panels F-I, control (black), *Flk2-cre.Cxcr4^fl/fl^* (red), *Pf4-cre.Cxcr4^fl/fl^* (green), *Lyz2-cre.Cxcr4^fl/fl^* (blue) and *Cd4-cre.Cxcr4^fl/fl^* (purple) mice. Bars indicate average, circles depict individual mice. n.s., not significant. P > 0.05; *, P < 0.05 by unpaired Student’s *t* test.

**Figure S5. Cytokine measurements, cKit signaling, and cKit blocking saturation.**

(A) Serum Thrombopoietin. (B) Soluble SCF in bone marrow interstitial fluid. (A and B) *Flk2-cre.Cxcr4^fl/+^* (black) and *Flk2-cre.Cxcr4^fl/fl^* (red) mice. (C) pSTAT3 and pSTAT5 staining in HSCs incubated with 600ng/ml mouse recombinant SCF (red) or medium (black). Numbers indicate GMFI. Bars indicate average, circles depict individual mice. *, P < 0.05 by unpaired Student’s *t* test. (D) cKit staining of bone marrow Lineage^-^ Sca-1^+^ cells in wild-type mice at day 3 after treatment with ACK2 (Rat-anti-mouse cKit) antibody or isotype control (200μg/mouse). Secondary antibody (anti-Rat IgG) staining of cells from isotype control treated mouse (gray), Secondary antibody staining of cells from ACK2 antibody treated mouse (red), ACK2 and secondary antibody staining of cells from ACK2 antibody treated mouse (blue).

## Notes

### Competing Interest Statement

The authors have declared no competing interest.

## References

Anders, S., P.T. Pyl, and W. Huber. 2015. HTSeq--a Python framework to work with high-throughput sequencing data. Bioinformatics 31:166–169.

Anderson, D.M., S.D. Lyman, A. Baird, J.M. Wignall, J. Eisenman, C. Rauch, C.J. March, H.S. Boswell, S.D. Gimpel, D. Cosman, and et al. 1990. Molecular cloning of mast cell growth factor, a hematopoietin that is active in both membrane bound and soluble forms. Cell 63:235–243.

Arojo, O.A., X. Ouyang, D. Liu, T. Meng, S.M. Kaech, J.P. Pereira, and B. Su. 2018. Active mTORC2 Signaling in Naive T Cells Suppresses Bone Marrow Homing by Inhibiting CXCR4 Expression. J Immunol 201:908–915.

Asada, N., Y. Kunisaki, H. Pierce, Z. Wang, N.F. Fernandez, A. Birbrair, A. Ma’ayan, and P.S. Frenette. 2017. Differential cytokine contributions of perivascular haematopoietic stem cell niches. Nat Cell Biol 19:214–223.

Baccin, C., J. Al-Sabah, L. Velten, P.M. Helbling, F. Grunschlager, P. Hernandez-Malmierca, C. Nombela-Arrieta, L.M. Steinmetz, A. Trumpp, and S. Haas. 2020. Combined single-cell and spatial transcriptomics reveal the molecular, cellular and spatial bone marrow niche organization. Nat Cell Biol 22:38–48.

Barker, J.E. 1994. Sl/Sld hematopoietic progenitors are deficient in situ. Exp Hematol 22:174–177.

Barker, J.E. 1997. Early transplantation to a normal microenvironment prevents the development of Steel hematopoietic stem cell defects. Exp Hematol 25:542–547.

Baryawno, N., D. Przybylski, M.S. Kowalczyk, Y. Kfoury, N. Severe, K. Gustafsson, K.D. Kokkaliaris, F. Mercier, M. Tabaka, M. Hofree, D. Dionne, A. Papazian, D. Lee, O. Ashenberg, A. Subramanian, E.D. Vaishnav, O. Rozenblatt-Rosen, A. Regev, and D.T. Scadden. 2019. A Cellular Taxonomy of the Bone Marrow Stroma in Homeostasis and Leukemia. Cell 177:1915–1932 e1916.

Beck, T.C., A.C. Gomes, J.G. Cyster, and J.P. Pereira. 2014. CXCR4 and a cell-extrinsic mechanism control immature B lymphocyte egress from bone marrow. The Journal of experimental medicine 211:2567–2581.

Boyer, S.W., A.V. Schroeder, S. Smith-Berdan, and E.C. Forsberg. 2011. All hematopoietic cells develop from hematopoietic stem cells through Flk2/Flt3-positive progenitor cells. Cell Stem Cell 9:64–73.

Bruns, I., D. Lucas, S. Pinho, J. Ahmed, M.P. Lambert, Y. Kunisaki, C. Scheiermann, L. Schiff, M. Poncz, A. Bergman, and P.S. Frenette. 2014. Megakaryocytes regulate hematopoietic stem cell quiescence through CXCL4 secretion. Nat Med 20:1315–1320.

Busch, K., K. Klapproth, M. Barile, M. Flossdorf, T. Holland-Letz, S.M. Schlenner, M. Reth, T. Hofer, and H.R. Rodewald. 2015. Fundamental properties of unperturbed haematopoiesis from stem cells in vivo. Nature 518:542–546.

Chaix, J., S.A. Nish, W.H. Lin, N.J. Rothman, L. Ding, E.J. Wherry, and S.L. Reiner. 2014. Cutting edge: CXCR4 is critical for CD8+ memory T cell homeostatic self-renewal but not rechallenge self-renewal. J Immunol 193:1013–1016.

Challen, G.A., N.C. Boles, S.M. Chambers, and M.A. Goodell. 2010. Distinct hematopoietic stem cell subtypes are differentially regulated by TGF-beta1. Cell Stem Cell 6:265–278.

Chhabra, A., A.M. Ring, K. Weiskopf, P.J. Schnorr, S. Gordon, A.C. Le, H.S. Kwon, N.G. Ring, J. Volkmer, P.Y. Ho, S. Tseng, I.L. Weissman, and J.A. Shizuru. 2016. Hematopoietic stem cell transplantation in immunocompetent hosts without radiation or chemotherapy. Sci Transl Med 8:351ra105.

Chow, A., D. Lucas, A. Hidalgo, S. Mendez-Ferrer, D. Hashimoto, C. Scheiermann, M. Battista, M. Leboeuf, C. Prophete, N. van Rooijen, M. Tanaka, M. Merad, and P.S. Frenette. 2011. Bone marrow CD169+ macrophages promote the retention of hematopoietic stem and progenitor cells in the mesenchymal stem cell niche. The Journal of experimental medicine 208:261–271.

Christodoulou, C., J.A. Spencer, S.A. Yeh, R. Turcotte, K.D. Kokkaliaris, R. Panero, A. Ramos, G. Guo, N. Seyedhassantehrani, T.V. Esipova, S.A. Vinogradov, S. Rudzinskas, Y. Zhang, A.S. Perkins, S.H. Orkin, R.A. Calogero, T. Schroeder, C.P. Lin, and F.D. Camargo. 2020. Live-animal imaging of native haematopoietic stem and progenitor cells. Nature 578:278–283.

Chung, Y.J., B.B. Park, Y.J. Kang, T.M. Kim, C.J. Eaves, and I.H. Oh. 2006. Unique effects of Stat3 on the early phase of hematopoietic stem cell regeneration. Blood 108:1208–1215.

Comazzetto, S., M.M. Murphy, S. Berto, E. Jeffery, Z. Zhao, and S.J. Morrison. 2019. Restricted Hematopoietic Progenitors and Erythropoiesis Require SCF from Leptin Receptor+ Niche Cells in the Bone Marrow. Cell Stem Cell 24:477–486 e476.

Cordeiro Gomes, A., T. Hara, V.Y. Lim, D. Herndler-Brandstetter, E. Nevius, T. Sugiyama, S. Tani-Ichi, S. Schlenner, E. Richie, H.R. Rodewald, R.A. Flavell, T. Nagasawa, K. Ikuta, and J.P. Pereira. 2016. Hematopoietic Stem Cell Niches Produce Lineage-Instructive Signals to Control Multipotent Progenitor Differentiation. Immunity 45:1219–1231.

Cyster, J.G., S.B. Hartley, and C.C. Goodnow. 1994. Competition for follicular niches excludes self-reactive cells from the recirculating B-cell repertoire. Nature 371:389–395.

Dance, A. 2019. Core Concept: Cells nibble one another via the under-appreciated process of trogocytosis. Proc Natl Acad Sci U S A 116:17608–17610.

Dann, E., N.C. Henderson, S.A. Teichmann, M.D. Morgan, and J.C. Marioni. 2020. <em>Milo:</em> differential abundance testing on single-cell data using k-NN graphs. bioRxiv 2020.2011.2023.393769.

Decker, M., J. Leslie, Q. Liu, and L. Ding. 2018. Hepatic thrombopoietin is required for bone marrow hematopoietic stem cell maintenance. Science (New York, N.Y 360:106–110.

Ding, L., and S.J. Morrison. 2013. Haematopoietic stem cells and early lymphoid progenitors occupy distinct bone marrow niches. Nature 495:231–235.

Ding, L., T.L. Saunders, G. Enikolopov, and S.J. Morrison. 2012. Endothelial and perivascular cells maintain haematopoietic stem cells. Nature 481:457–462.

Finak, G., A. McDavid, M. Yajima, J. Deng, V. Gersuk, A.K. Shalek, C.K. Slichter, H.W. Miller, M.J. McElrath, M. Prlic, P.S. Linsley, and R. Gottardo. 2015. MAST: a flexible statistical framework for assessing transcriptional changes and characterizing heterogeneity in single-cell RNA sequencing data. Genome Biol 16:278.

Fistonich, C., S. Zehentmeier, J.J. Bednarski, R. Miao, H. Schjerven, B.P. Sleckman, and J.P. Pereira. 2018. Cell circuits between B cell progenitors and IL-7(+) mesenchymal progenitor cells control B cell development. J Exp Med 215:2586–2599.

Gekas, C., and T. Graf. 2013. CD41 expression marks myeloid-biased adult hematopoietic stem cells and increases with age. Blood 121:4463–4472.

Greenbaum, A., Y.M. Hsu, R.B. Day, L.G. Schuettpelz, M.J. Christopher, J.N. Borgerding, T. Nagasawa, and D.C. Link. 2013. CXCL12 in early mesenchymal progenitors is required for haematopoietic stem-cell maintenance. Nature 495:227–230.

Hafemeister, C., and R. Satija. 2019. Normalization and variance stabilization of single-cell RNA-seq data using regularized negative binomial regression. Genome Biol 20:296.

Hirata, Y., K. Furuhashi, H. Ishii, H.W. Li, S. Pinho, L. Ding, S.C. Robson, P.S. Frenette, and J. Fujisaki. 2018. CD150(high) Bone Marrow Tregs Maintain Hematopoietic Stem Cell Quiescence and Immune Privilege via Adenosine. Cell Stem Cell 22:445–453 e445.

Joly, E., and D. Hudrisier. 2003. What is trogocytosis and what is its purpose? Nature immunology 4:815.

Kim, T.S., and E.C. Shin. 2019. The activation of bystander CD8(+) T cells and their roles in viral infection. Exp Mol Med 51:1–9.

Li, H., B. Handsaker, A. Wysoker, T. Fennell, J. Ruan, N. Homer, G. Marth, G. Abecasis, R. Durbin, and S. Genome Project Data Processing. 2009. The Sequence Alignment/Map format and SAMtools. Bioinformatics 25:2078–2079.

Lim, V.Y., S. Zehentmeier, C. Fistonich, and J.P. Pereira. 2017. A Chemoattractant-Guided Walk Through Lymphopoiesis: From Hematopoietic Stem Cells to Mature B Lymphocytes. Adv Immunol 134:47–88.

Linnekin, D. 1999. Early signaling pathways activated by c-Kit in hematopoietic cells. Int J Biochem Cell Biol 31:1053–1074.

Lo Celso, C., H.E. Fleming, J.W. Wu, C.X. Zhao, S. Miake-Lye, J. Fujisaki, D. Cote, D.W. Rowe, C.P. Lin, and D.T. Scadden. 2009. Live-animal tracking of individual haematopoietic stem/progenitor cells in their niche. Nature 457:92–96.

Love, M.I., W. Huber, and S. Anders. 2014. Moderated estimation of fold change and dispersion for RNA-seq data with DESeq2. Genome Biol 15:550.

McInnes, L.H., J.; Saul, N.; Großberger, L. 2018. UMAP: Uniform Manifold Approximation and Projection.. Journal of Open Source Software 3:861.

Mendez-Ferrer, S., D. Bonnet, D.P. Steensma, R.P. Hasserjian, I.M. Ghobrial, J.G. Gribben, M. Andreeff, and D.S. Krause. 2020. Bone marrow niches in haematological malignancies. Nat Rev Cancer 20:285–298.

Mendez-Ferrer, S., T.V. Michurina, F. Ferraro, A.R. Mazloom, B.D. Macarthur, S.A. Lira, D.T. Scadden, A. Ma’ayan, G.N. Enikolopov, and P.S. Frenette. 2010. Mesenchymal and haematopoietic stem cells form a unique bone marrow niche. Nature 466:829–834.

Miao, R., V.Y. Lim, N. Kothapalli, Y. Ma, J. Fossati, S. Zehentmeier, R. Sun, and J.P. Pereira. 2020. Hematopoietic Stem Cell Niches and Signals Controlling Immune Cell Development and Maintenance of Immunological Memory. Front Immunol 11:600127.

Morrison, S.J., and D.T. Scadden. 2014. The bone marrow niche for haematopoietic stem cells. Nature 505:327–334.

Omatsu, Y., T. Sugiyama, H. Kohara, G. Kondoh, N. Fujii, K. Kohno, and T. Nagasawa. 2010. The Essential Functions of Adipo-osteogenic Progenitors as the Hematopoietic Stem and Progenitor Cell Niche. Immunity 33:387–399.

Pertea, M., D. Kim, G.M. Pertea, J.T. Leek, and S.L. Salzberg. 2016. Transcript-level expression analysis of RNA-seq experiments with HISAT, StringTie and Ballgown. Nat Protoc 11:1650–1667.

Pietras, E.M., D. Reynaud, Y.A. Kang, D. Carlin, F.J. Calero-Nieto, A.D. Leavitt, J.M. Stuart, B. Gottgens, and E. Passegue. 2015. Functionally Distinct Subsets of Lineage-Biased Multipotent Progenitors Control Blood Production in Normal and Regenerative Conditions. Cell Stem Cell 17:35–46.

Sawai, Catherine M., S. Babovic, S. Upadhaya, David J.H.F. Knapp, Y. Lavin, Colleen M. Lau, A. Goloborodko, J. Feng, J. Fujisaki, L. Ding, Leonid A. Mirny, M. Merad, Connie J. Eaves, and B. Reizis. 2016. Hematopoietic Stem Cells Are the Major Source of Multilineage Hematopoiesis in Adult Animals. Immunity

Severe, N., N.M. Karabacak, K. Gustafsson, N. Baryawno, G. Courties, Y. Kfoury, K.D. Kokkaliaris, C. Rhee, D. Lee, E.W. Scadden, J.E. Garcia-Robledo, T. Brouse, M. Nahrendorf, M. Toner, and D.T. Scadden. 2019. Stress-Induced Changes in Bone Marrow Stromal Cell Populations Revealed through Single-Cell Protein Expression Mapping. Cell Stem Cell 25:570–583 e577.

Shen, B., A. Tasdogan, J.M. Ubellacker, J. Zhang, E.D. Nosyreva, L. Du, M.M. Murphy, S. Hu, Y. Yi, N. Kara, X. Liu, S. Guela, Y. Jia, V. Ramesh, C. Embree, E.C. Mitchell, Y.C. Zhao, L.A. Ju, Z. Hu, G.M. Crane, Z. Zhao, R. Syeda, and S.J. Morrison. 2021. A mechanosensitive peri-arteriolar niche for osteogenesis and lymphopoiesis. Nature

Shimoto, M., T. Sugiyama, and T. Nagasawa. 2017. Numerous niches for hematopoietic stem cells remain empty during homeostasis. Blood 129:2124–2131.

Silberstein, L., K.A. Goncalves, P.V. Kharchenko, R. Turcotte, Y. Kfoury, F. Mercier, N. Baryawno, N. Severe, J. Bachand, J.A. Spencer, A. Papazian, D. Lee, B.R. Chitteti, E.F. Srour, J. Hoggatt, T. Tate, C. Lo Celso, N. Ono, S. Nutt, J. Heino, K. Sipila, T. Shioda, M. Osawa, C.P. Lin, G.F. Hu, and D.T. Scadden. 2016. Proximity-Based Differential Single-Cell Analysis of the Niche to Identify Stem/Progenitor Cell Regulators. Cell Stem Cell

Stuart, T., A. Butler, P. Hoffman, C. Hafemeister, E. Papalexi, W.M. Mauck, 3rd, Y. Hao, M. Stoeckius, P. Smibert, and R. Satija. 2019. Comprehensive Integration of Single-Cell Data. Cell 177:1888–1902 e1821.

Sugiyama, T., H. Kohara, M. Noda, and T. Nagasawa. 2006. Maintenance of the hematopoietic stem cell pool by CXCL12-CXCR4 chemokine signaling in bone marrow stromal cell niches. Immunity 25:977–988.

Tikhonova, A.N., I. Dolgalev, H. Hu, K.K. Sivaraj, E. Hoxha, A. Cuesta-Dominguez, S. Pinho, I. Akhmetzyanova, J. Gao, M. Witkowski, M. Guillamot, M.C. Gutkin, Y. Zhang, C. Marier, C. Diefenbach, S. Kousteni, A. Heguy, H. Zhong, D.R. Fooksman, J.M. Butler, A. Economides, P.S. Frenette, R.H. Adams, R. Satija, A. Tsirigos, and I. Aifantis. 2019. The bone marrow microenvironment at single-cell resolution. Nature 569:222–228.

Tokoyoda, K., S. Zehentmeier, A.N. Hegazy, I. Albrecht, J.R. Grun, M. Lohning, and A. Radbruch. 2009. Professional memory CD4+ T lymphocytes preferentially reside and rest in the bone marrow. Immunity 30:721–730.

Tullai, J.W., M.E. Schaffer, S. Mullenbrock, G. Sholder, S. Kasif, and G.M. Cooper. 2007. Immediate-early and delayed primary response genes are distinct in function and genomic architecture. The Journal of biological chemistry 282:23981–23995.

Upadhaya, S., O. Krichevsky, I. Akhmetzyanova, C.M. Sawai, D.R. Fooksman, and B. Reizis. 2020. Intravital Imaging Reveals Motility of Adult Hematopoietic Stem Cells in the Bone Marrow Niche. Cell Stem Cell 27:336–345 e334.

van Dijk, D., R. Sharma, J. Nainys, K. Yim, P. Kathail, A.J. Carr, C. Burdziak, K.R. Moon, C.L. Chaffer, D. Pattabiraman, B. Bierie, L. Mazutis, G. Wolf, S. Krishnaswamy, and D. Pe’er. 2018. Recovering Gene Interactions from Single-Cell Data Using Data Diffusion. Cell 174:716–729 e727.

Victora, G.D., and M.C. Nussenzweig. 2012. Germinal centers. Annual review of immunology 30:429–457.

Wei, Q., and P.S. Frenette. 2018. Niches for Hematopoietic Stem Cells and Their Progeny. Immunity 48:632–648.

Xu, C., X. Gao, Q. Wei, F. Nakahara, S.E. Zimmerman, J. Mar, and P.S. Frenette. 2018. Stem cell factor is selectively secreted by arterial endothelial cells in bone marrow. Nat Commun 9:2449.

Zehentmeier, S., and J.P. Pereira. 2019. Cell circuits and niches controlling B cell development. Immunol Rev 289:142–157.

Zhang, J., Q. Wu, C.B. Johnson, G. Pham, J.M. Kinder, A. Olsson, A. Slaughter, M. May, B. Weinhaus, A. D’Alessandro, J.D. Engel, J.X. Jiang, J.M. Kofron, L.F. Huang, V.B.S. Prasath, S.S. Way, N. Salomonis, H.L. Grimes, and D. Lucas. 2021. In situ mapping identifies distinct vascular niches for myelopoiesis. Nature 590:457–462.

Zhao, M., J.M. Perry, H. Marshall, A. Venkatraman, P. Qian, X.C. He, J. Ahamed, and L. Li. 2014. Megakaryocytes maintain homeostatic quiescence and promote post-injury regeneration of hematopoietic stem cells. Nat Med 20:1321–1326.

Zheng, G.X., J.M. Terry, P. Belgrader, P. Ryvkin, Z.W. Bent, R. Wilson, S.B. Ziraldo, T.D. Wheeler, G.P. McDermott, J. Zhu, M.T. Gregory, J. Shuga, L. Montesclaros, J.G. Underwood, D.A. Masquelier, S.Y. Nishimura, M. Schnall-Levin, P.W. Wyatt, C.M. Hindson, R. Bharadwaj, A. Wong, K.D. Ness, L.W. Beppu, H.J. Deeg, C. McFarland, K.R. Loeb, W.J. Valente, N.G. Ericson, E.A. Stevens, J.P. Radich, T.S. Mikkelsen, B.J. Hindson, and J.H. Bielas. 2017. Massively parallel digital transcriptional profiling of single cells. Nat Commun 8:14049.

