## Supplementary figures and images for "Competition between hematopoietic stem and progenitor cells controls hematopoietic stem cell compartment size"

### Supplemental Figure 1

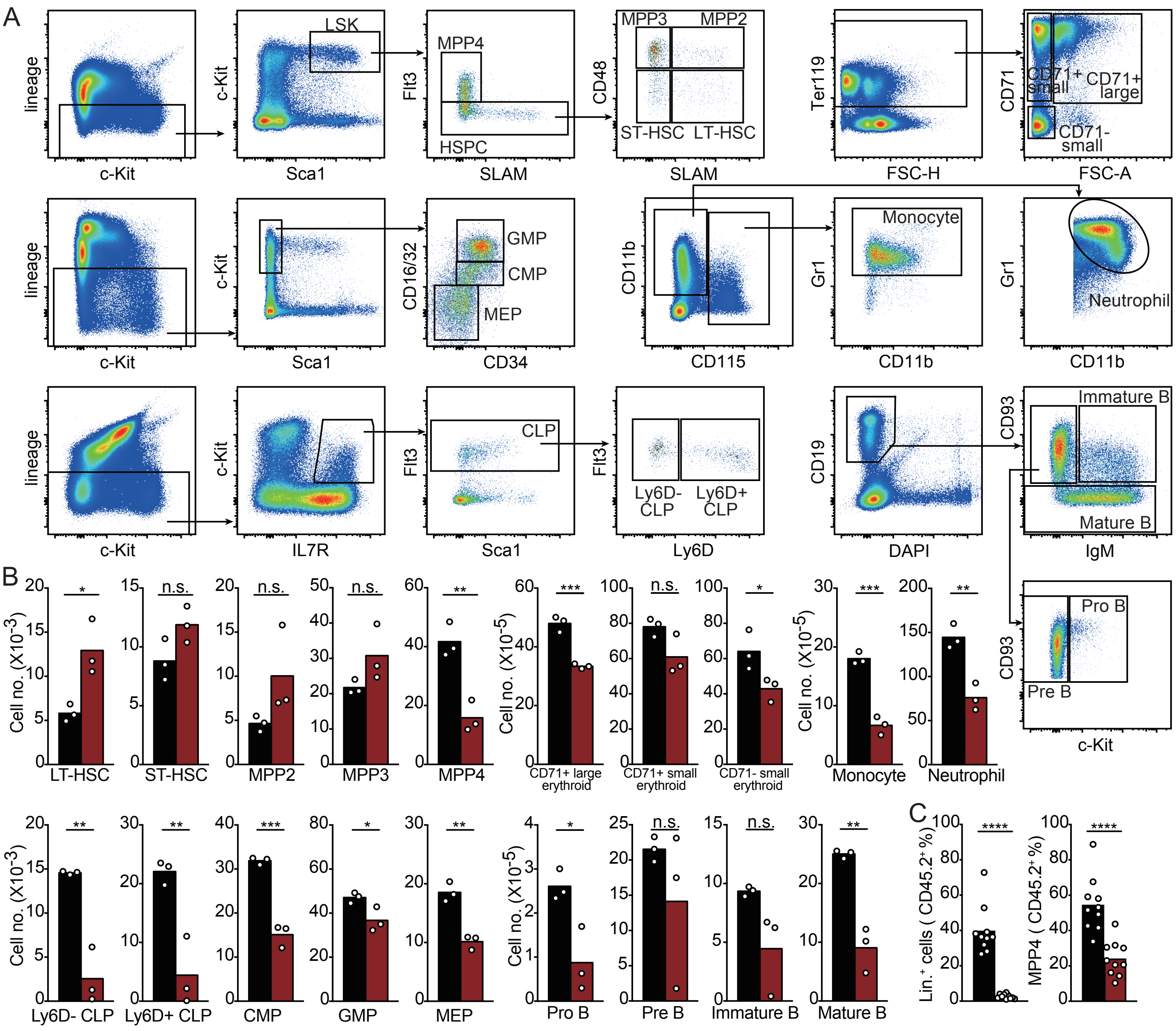


Supplementary Figure 1.

### Supplemental Figure 2

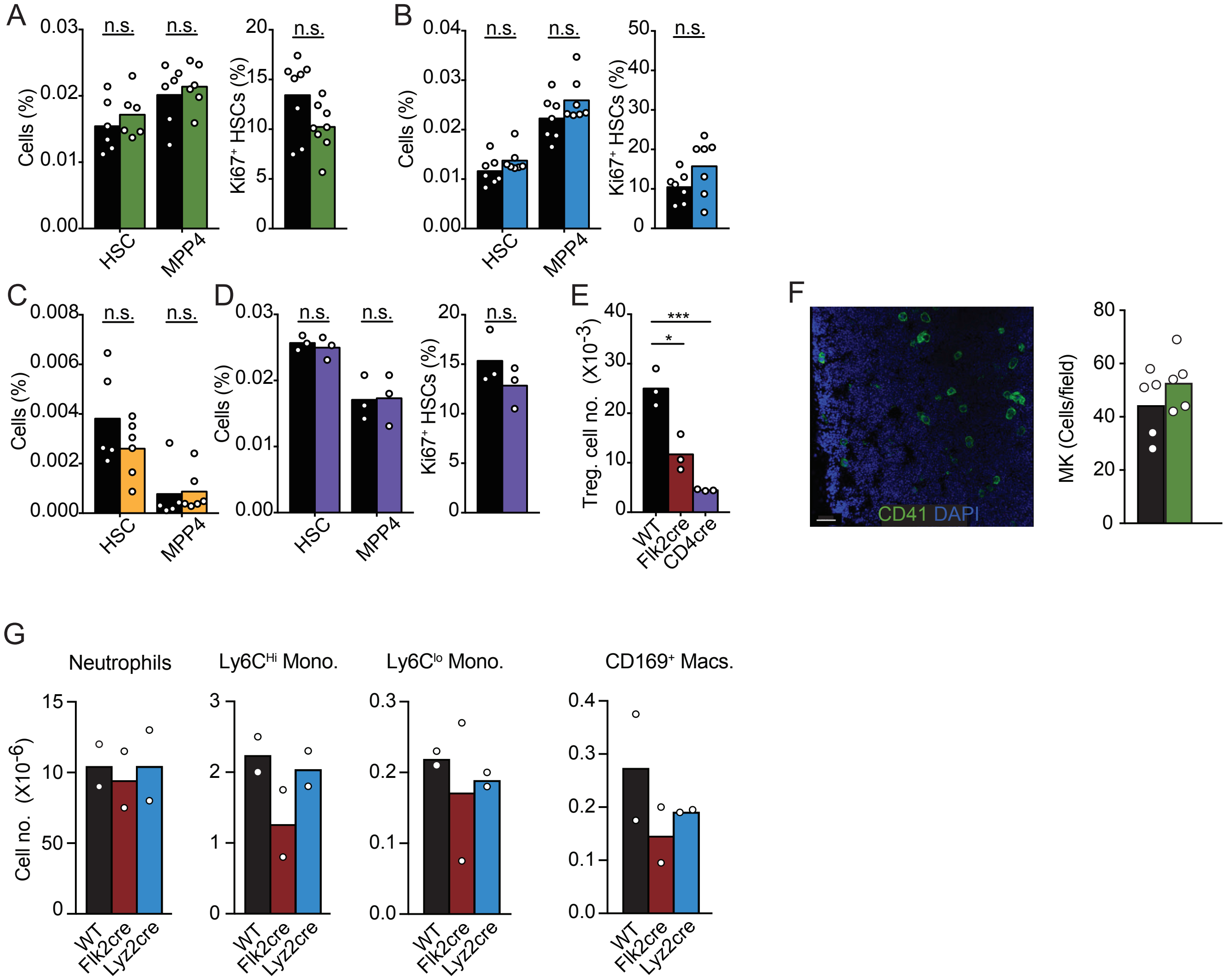


Supplementary Figure 2.

### Supplemental Figure 3

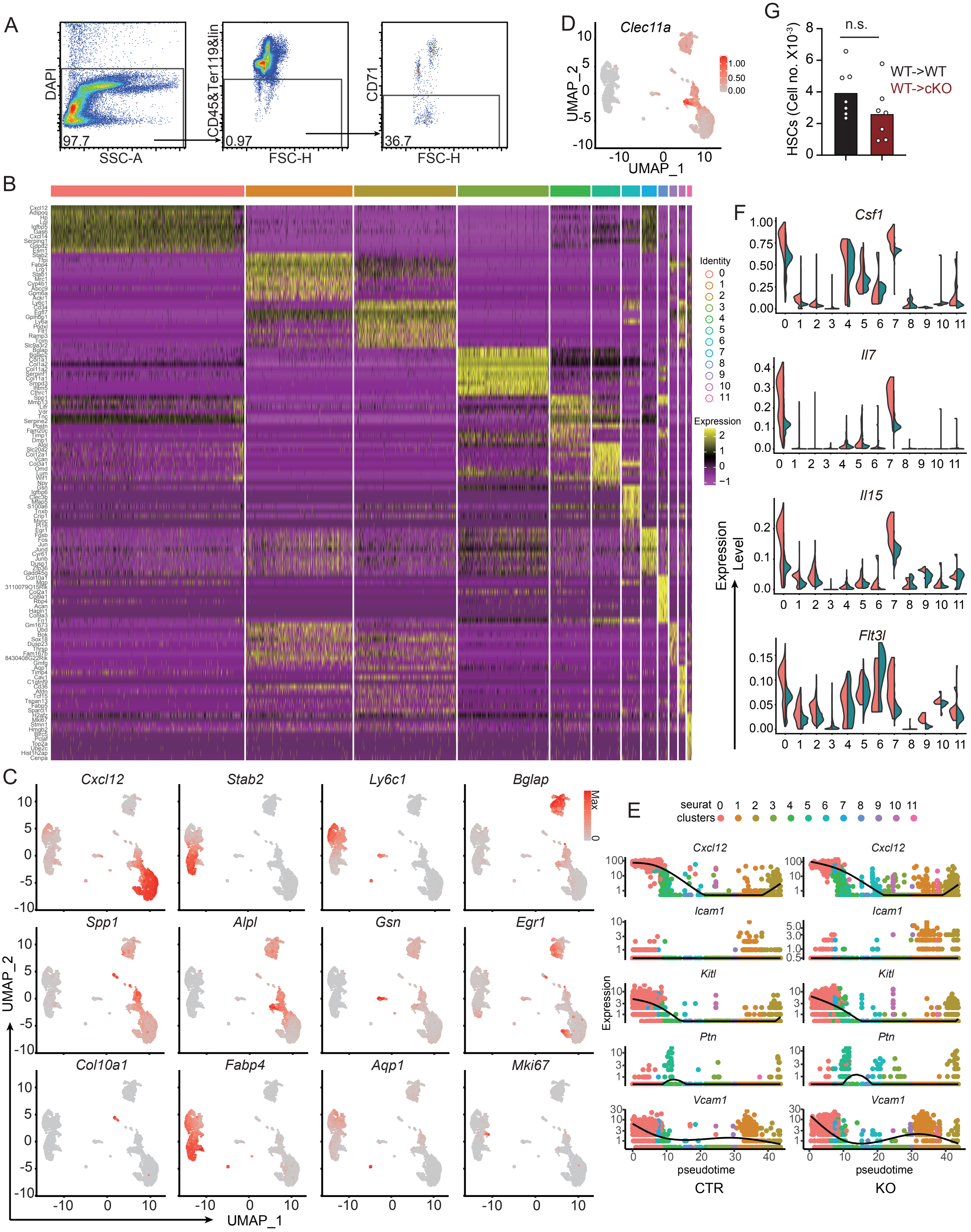


Supplementary Figure 3.

### Supplemental Figure 4

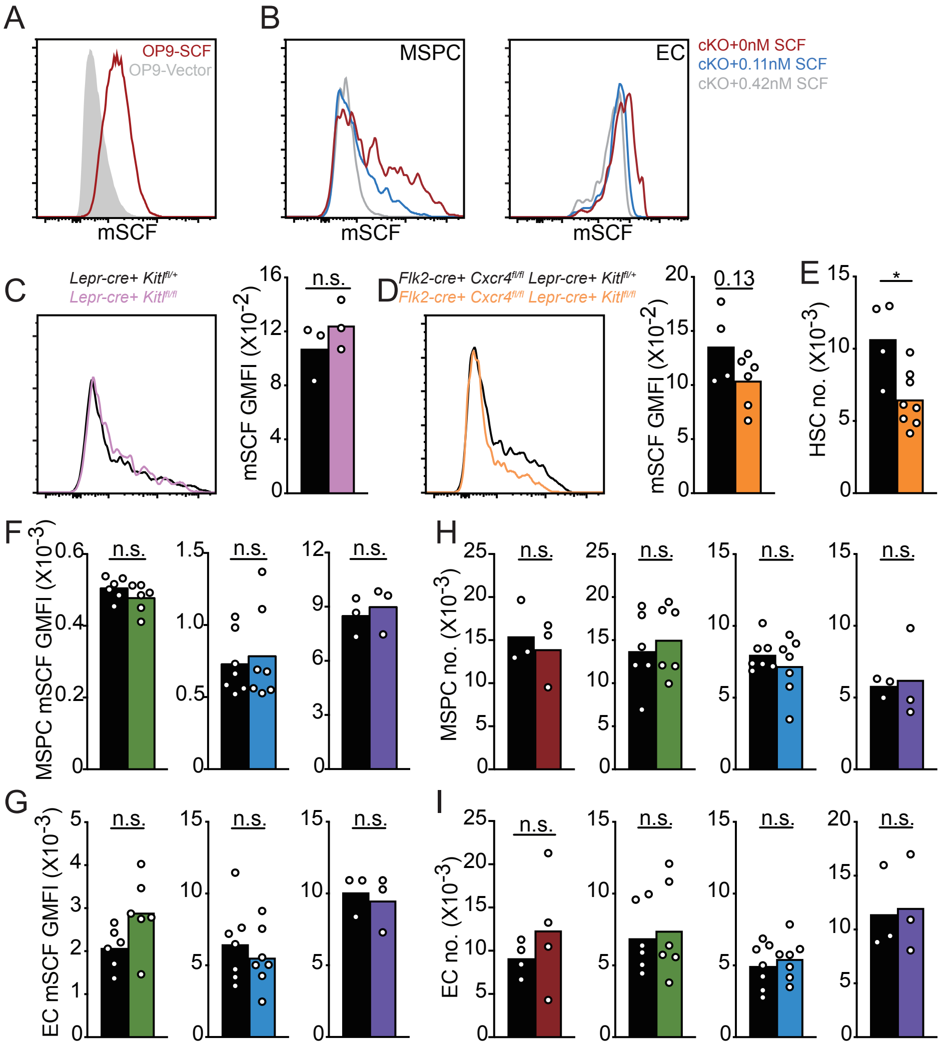


Supplementary Figure 4.

### Supplemental Figure 5

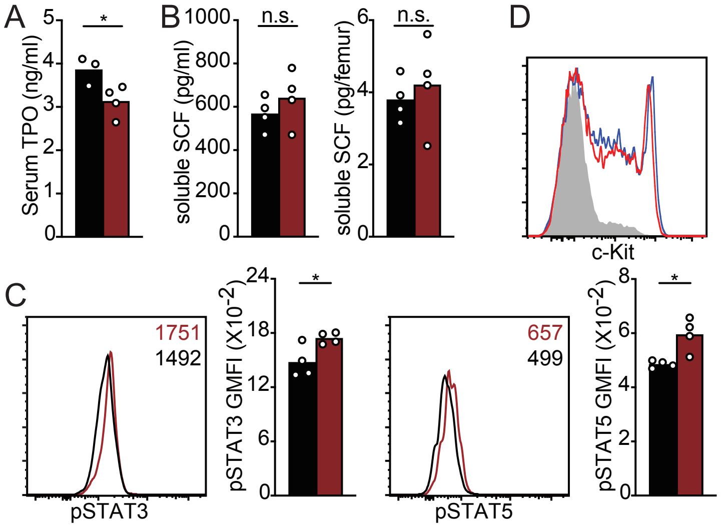


Supplementary Figure 5.
